# Interactions between the picornavirus 3C(D) main protease and RNA induce liquid-liquid phase separation

**DOI:** 10.1101/2025.04.14.648818

**Authors:** Somnath Mondal, Saumyak Mukherjee, Kevin E.W. Namitz, Neela Yennawar, David D Boehr

## Abstract

The picornavirus 3CD protein is a precursor to the 3C main protease and the 3D RNA-dependent RNA polymerase. In addition to its functions in proteolytic processing of the virus polyprotein and cleavage of key host factors, the 3C domain interacts with cis-acting replication elements (CREs) within the viral genome to regulate replication and translation events. We investigated the molecular determinants of RNA binding to 3C using a wide range of biophysical and computational methods. These studies showed that 3C binds to a broad spectrum of RNA oligonucleotides, displaying minimal dependence on RNA sequence and structure. However, they also uncovered a novel aspect of these interactions, that is, 3C-RNA binding can induce liquid-liquid phase separation (LLPS), with 3CD-RNA interactions likewise leading to LLPS. This may be a general phenomenon for other 3C and 3C-like proteases, and polyproteins incorporating 3C domains. These findings have potential implications in understanding virally induced apoptosis and controlling stress granules, which involve LLPS and include other proteins with known interactions with 3C/3CD.

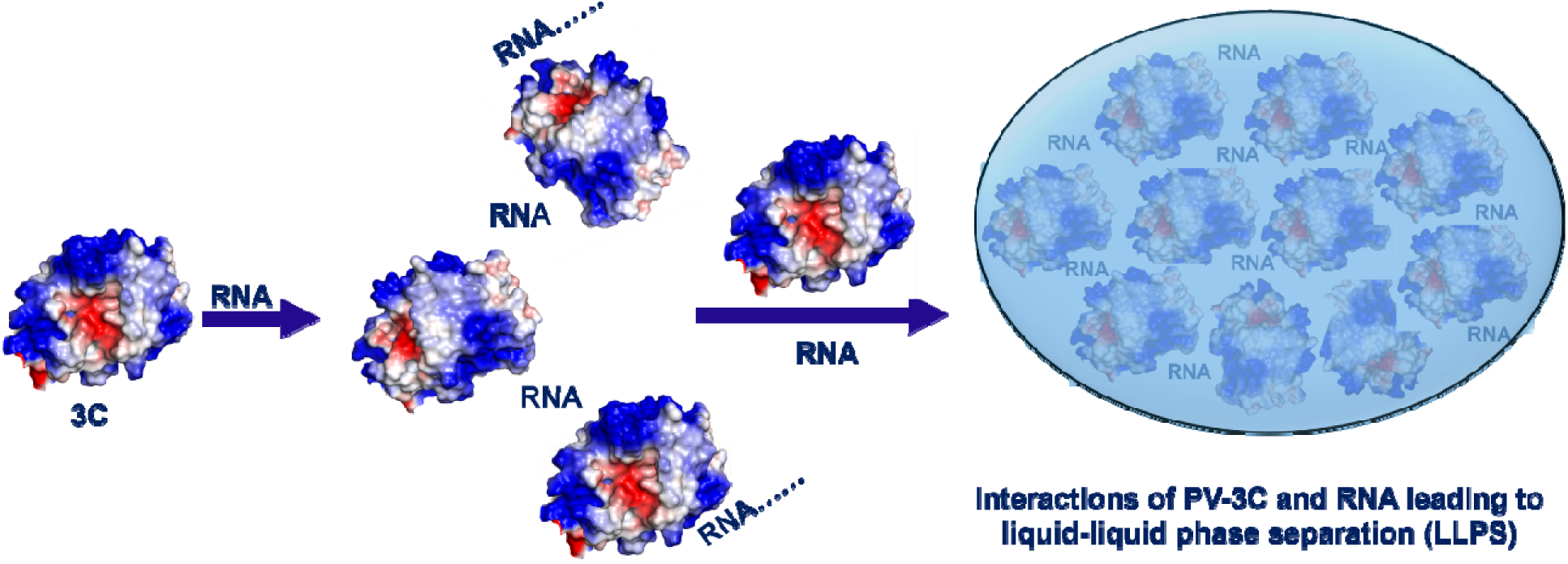

## INTRODUCTION

Positive-strand RNA viruses generate a polyprotein that must then be proteolytically processed into the components necessary for virus replication and packaging^1,2^. These viruses have evolved strategies to maximize their genomic information content. For example, intermediates in the proteolytic processing pathway may have different and/or emergent functions compared to their fully processed counterparts. As well, many viral proteins have multiple functions. These ideas are exemplified by the picornavirus 3CD, 3C and 3D proteins^3–5^. 3CD is comprised of domains encompassing the 3C protease and the 3D RNA-dependent RNA polymerase^3,4,6^. The 3CD and/or 3C proteases cleave the viral polyprotein and host cell defence proteins targeting viral RNA synthesis and replication^7–9^. The 3C domain also interacts with cis-acting replication elements (CREs; oriL, oriI, and oriR) to regulate viral replication and translation, helping to coordinate the virus life cycle^10,11^. The molecular determinants governing 3C-RNA interactions and how these interactions may affect other 3C/3CD functions are poorly understood.

3C and 3C-like proteases are found in a wide range of positive-strand RNA viruses, particularly picornaviruses (poliovirus, PV; human rhinovirus, HRV; hepatitis A virus, HAV; foot-and-mouth disease virus, FMDV; coxsackievirus B, CVB; cardio virus; enterovirus-D68, EV-D68; and enterovirus 71, EV71), coronaviruses (severe acute respiratory syndrome coronavirus 1 and 2, SARS-CoV and SARS-CoV-2; Middle East respiratory syndrome coronavirus, MERS-CoV), alphaviruses (Chikungunya virus, eastern equine encephalitis virus), flaviviruses (dengue virus, zika virus, West Nile virus, yellow fever virus), caliciviruses (norovirus, Sapo virus), and astroviruses^11–14^. The 3C three-dimensional structures are highly conserved, consisting of two antiparallel, six-stranded β-barrel domains, and catalytic mechanisms generally identical, resembling that of trypsin-like Ser proteases but with a Cys nucleophile (**Fig. 1**)^3,11,15^. In PV, the 3C catalytic triad consists of His40, Glu71, and Cys147, which are spatially arranged to facilitate the catalytic reaction.

**Figure 1.**
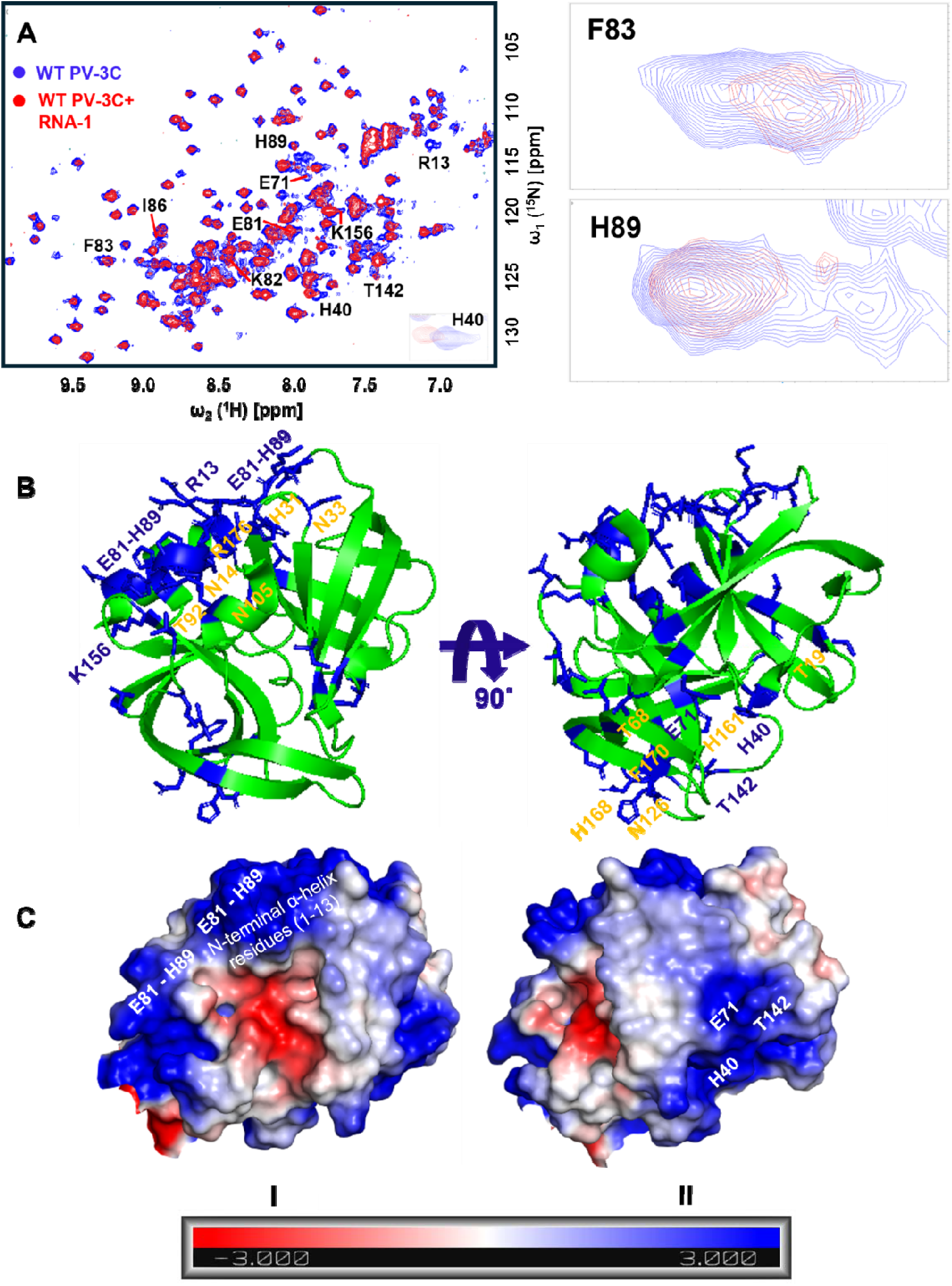
RNA binding leads to chemical shift changes for two oppositely faced residue clusters in PV-3C. (A) ^1^H-^15^N SOFAST-HMQC NMR spectra were compared with free wild-type PV-3C (blue) and RNA-1 (GGCGGCGUACUCCGG) bound PV-3C (red) in 1:0.5 stoichiometry; the residues with substantial chemical shift perturbation (CSP) are labelled in the spectra; zoomed in resonances represents CSP change for the corresponding residues. (B) Residues showing substantial CSPs in the presence of RNA are represented in blue on the PV-3C X-ray crystal structure (PDB ID: 1L1N). Note that the R13 peak was missing after the addition of RNA. Orange colored residues represent substantial CSPs in the presence of RNA-3-RNA-15 overall on PV-3C (see **Fig. S2-S14**). NMR CSPs were calculated using the following equation Δδ_combined =_ (Δδ_H_^2^ + (Δδ_N_/5)^2^)^0.5^, where Δδ_H_ and Δδ_N_ are the chemical shift differences between 3C with and without RNA for the backbone amide proton and nitrogen respectively; those with substantial CSPs have Δδ_combined_ greater than 0.02 ppm. The PV-3C protein concentration was 25 µM and RNA was added following a 1:0.5 stoichiometry using a buffer containing 10 mM HEPES pH 7.5 and 50 mM NaCl. The ^1^H-^15^N SOFAST-HMQC NMR experiment was recorded in a 600 MHz Bruker NEO spectrometer equipped with z-gradient triple resonance (^1^H, ^13^C, ^15^N) TCI cryogenic probe at 25_°_C. (C) The electrostatic surface potential map for PV-3C, highlighting Cluster-I and Cluster-II regions, with blue and red representing positively- and negatively-charged regions, respectively.

The 3C protein and/or the 3C domain in 3CD also have important RNA-binding capabilities. For example, 3CD interacts with the stem-loop d of oriL, enhancing the binding of the poly(rC)-binding protein 2 (PCBP2) to the stem-loop b^7,8,16^. Additionally, the RNA-binding abilities of both 3C and 3CD through oriI, are indispensable for the effective uridylylation of VPg protein^17^. Previous studies have identified the 80’s region of 3C as highly important for binding to the stem-loop of oriI^17^. Given the sequence/structural diversity of these RNA elements, it is unclear what factors determine these interactions. A fuller understanding of these interactions may provide new insights into the development of anti-viral therapeutics that disrupt these crucial interactions.

To better define the molecular determinants of RNA interactions with PV-3C and PV-3CD, we assessed the binding of a variety of RNA oligonucleotides with diverse sequences and secondary structure capabilities using a range of experimental and computational methods. While little specificity was found, these interactions surprisingly lead to higher-order complex formation and liquid-liquid phase separation (LLPS) for both PV-3C and PV-3CD. LLPS has previously been found in various viral infections, and it may play a role in virally induced apoptosis and virus dissemination through the formation/deformation of stress granules^18–20^. As such, these findings provide a potential new role for 3C/3CD in these processes and potentially reveal another level of regulatory processes involving these proteins.

## RESULTS

### PV-3C binds to a diverse set of RNA oligonucleotides

To better understand the molecular determinants of RNA binding to 3C, we collected NMR spectra for PV-3C in the presence and absence of 15 sequence- and structure-divergent RNA oligonucleotides (see **Fig. 1 and Table S1**). Some of these RNA oligonucleotides were based on sequences derived from oriL and/or oriR (e.g. RNA-1, 5’-GGCGGCGUACUCCGG-3’ from oriL in **Fig. 1 and Fig. S1**). Here, we focus on NMR spectra with RNA-1, but, notably, similar results were obtained for all RNA tested from RNA-2 to RNA-15 (**Fig. S2-S14**), suggesting that there is little to no sequence dependence of RNA binding to 3C using this set of RNA oligonucleotides.

[^15^N,^1^H] SOFAST-HMQC NMR spectra were collected after titrating ^15^N-labeled PV-3C with increasing concentrations of RNA-1^21^. The addition of RNA (shown in **Fig. 1**) led to chemical shift perturbations (CSPs) for ^1^H-^15^N backbone amide resonances belonging to amino acid residues in the previously identified RNA binding region (E81-H89), the N-terminal α residues (1-13), the active site residues (H40 and E71, part of the catalytic triad) and nearby residues T142 and K156 (**Fig. 1A**). These amino acid residues appear to belong to two clusters (**Fig. 1B**), Cluster-I and Cluster-II. Cluster-I is located near the KFRDI motif, which has been linked to RNA binding in earlier NMR investigations^17,22^ and mutational studies^23,24^. The N-terminal *h*1 helix is near this RNA binding region and has also been previously implicated in RNA binding^25^. Recent reports indicate that Coxsackievirus 3C protein interacts with cloverleaf stem-loop D RNA through the KFRDI motif, the N-terminal *h*1-helix, and K156 residues, which is consistent with Cluster-I residues observed in our NMR studies^26^. Cluster-II is on the opposite side of the protein, including the catalytic and adjacent residues (**Fig. 1B**). It is noted that we took into consideration the results for all the different RNA oligonucleotides (also see **Fig. S2-S14**) when identifying these two clusters. Intriguingly, mapping of electrostatic surface potentials indicates that Cluster-I and Cluster-II regions are positively charged regions, suggesting a mechanism by which both Clusters could interact with negatively charged RNA (**Fig. 1C**).

It is also noteworthy that further addition of RNA beyond 1:0.5 stoichiometry of 3C: RNA resulted in the disappearance of peaks in a nonspecific manner (**Fig. S16)**. While this complicates analysis and prevents assessment of binding affinity by NMR, the overall decrease in NMR signals would be consistent with the formation of larger 3C-RNA complexes in an RNA-independent manner.

### Small-angle X-ray scattering is consistent with RNA binding to the Cluster I region

Small-angle X-ray scattering (SAXS) experiments were also performed on a 1:1 3C: RNA-1 sample after the addition of the RNA-1 sample to 3C (at 25 µM concentration each as utilised for NMR, **Fig. 1**). Scattering curves and derived Guinear fits, Kratky plots and pair distance distribution (P(r)) functions provide important structural insights into the complex formation by identifying the size, shape, flexibility and distance changes on complexation (**Fig. 2, Fig S15**). The radii of gyration, maximum dimension of the particle and Porod volume all increases in the 3C-RNA complex (19.4Å, 60Å and 18901Å^3^ respectively) as compared to 3C alone (16.3Å, 48Å and 16531Å^3^ respectively) (**Fig. 2**). 3C shows a more compact envelope which elongates on addition of RNA, with the overall conformation of 3C not appearing to change substantially after the addition of RNA (**Fig. 2**).

**Figure 2.**
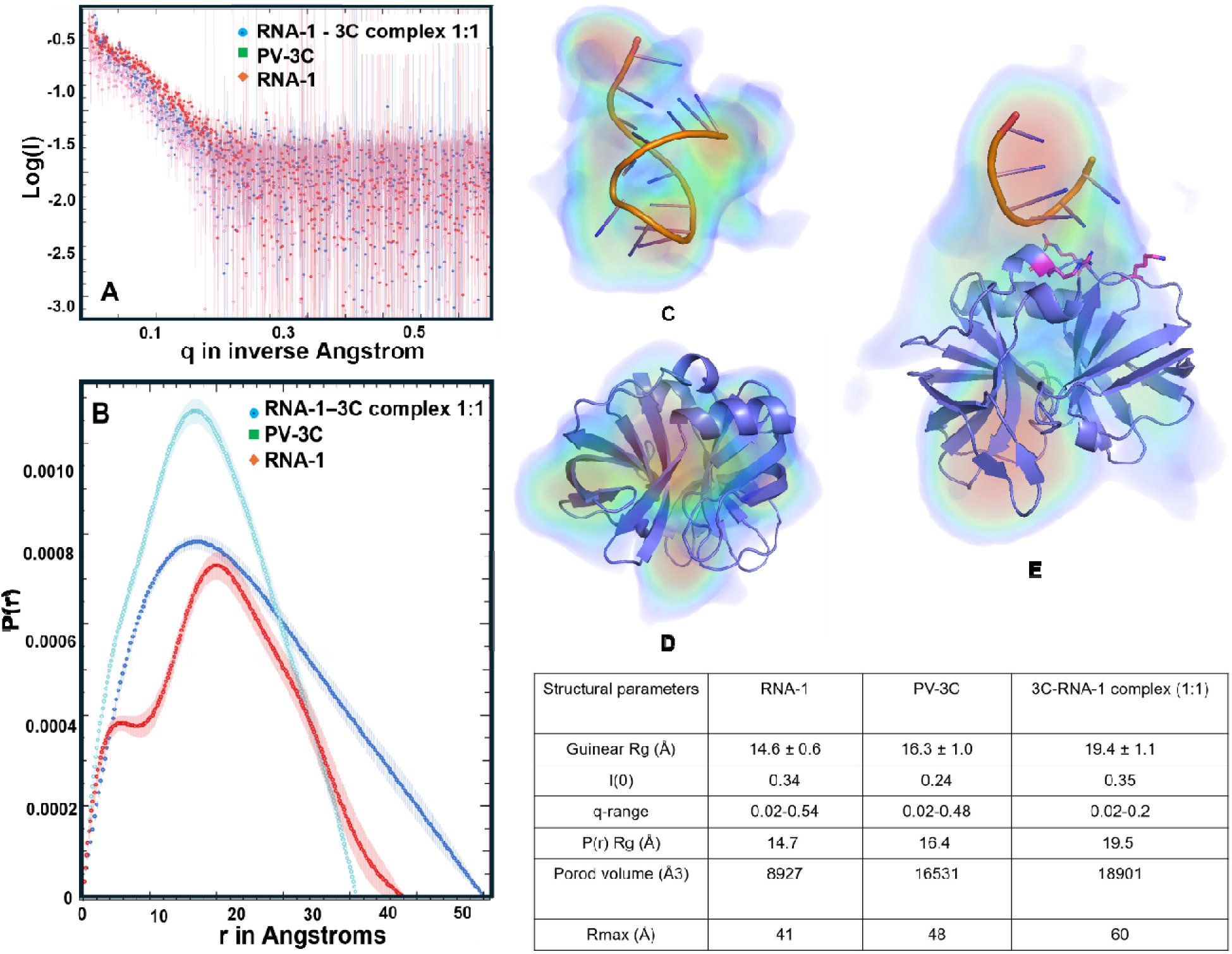
Small-angle X-ray scattering is consistent with RNA binding to the Cluster-I region of PV-3C. Small-angle X-ray scattering (SAXS) data of PV-3C, RNA-1, and 3C-RNA complex: (A) SAXS raw data overlay for PV-3C protein (square), RNA-1 (diamond), and 3C-RNA complex (sphere). (B) Pair distance distribution function, P(r) function represents the histogram of distances between pairs of points within the particle. P(r) for the 3C SAXS data sets peaking at 20.0 Å in green, and 25 Å in 3C+RNA-1 in blue. (C), (D), and (E) demonstrate a solvent envelope fitting a monomer model of RNA-1, PV-3C, and 3C-RNA complex from the DENSS electron density map and PyMOL. We are showing only the interacting regions of RNA-1 as the stem-loop is not visible in the DENSS map. Pink residues in PV-3C comes from Cluster-I, identified from the NMR experiments. Structural parameters of 3C, RNA-1, and 3C-RNA-1 complex (1:1) are presented in the table.

The Direct Evaluation of Non-Solvent Scattering (DENSS) method in SAXS extracts extensive structural information about samples by excluding the scattering contributions from the solvent and the solute. Solvent envelopes were generated from DENSS for apo and complex states. The solvent envelope for RNA-1 alone (**Fig. 2C**) and 3C protein alone (**Fig. 2D**) matched the monomeric models for the respective species. The 3C-RNA complex (**Fig. 2E**) had an envelope that fit with a 1:1 3C:RNA complex stoichiometry with a chi-square fit of 1.9 as from Crysol (**Fig. S15**), although missing part of density for the terminal region of the RNA, likely owing to its flexibility. Moreover, the model for the 3C-RNA complex is consistent with the RNA binding to the Cluster-I region of 3C (**Fig. 2E**).

### AUC reveals multimeric complex formation between PV-3C and RNA

NMR experiments with higher concentrations of RNA led to an overall decrease in peak intensities (**Fig. S16**), which might be explained by the formation of multimeric complexes with 3C and RNA. To test this possibility, we utilized Sedimentation Velocity Analytical Ultracentrifugation (SV-AUC), which provides information about the oligomerization of macromolecules in solutions. In the absence of RNA, PV-3C is almost entirely monomeric (**Fig. S17**). At a 1:1 molar ratio of PV-3C and RNA-1, there is only evidence of a single species, which is consistent with the SAXS results (**Figs. 2, 3**). However, at higher ratios of RNA (i.e. 1:2 3C: RNA-1 and above), various multimeric complexes appear likely owing to oligomerization (**Fig. 3**).

**Figure 3.**
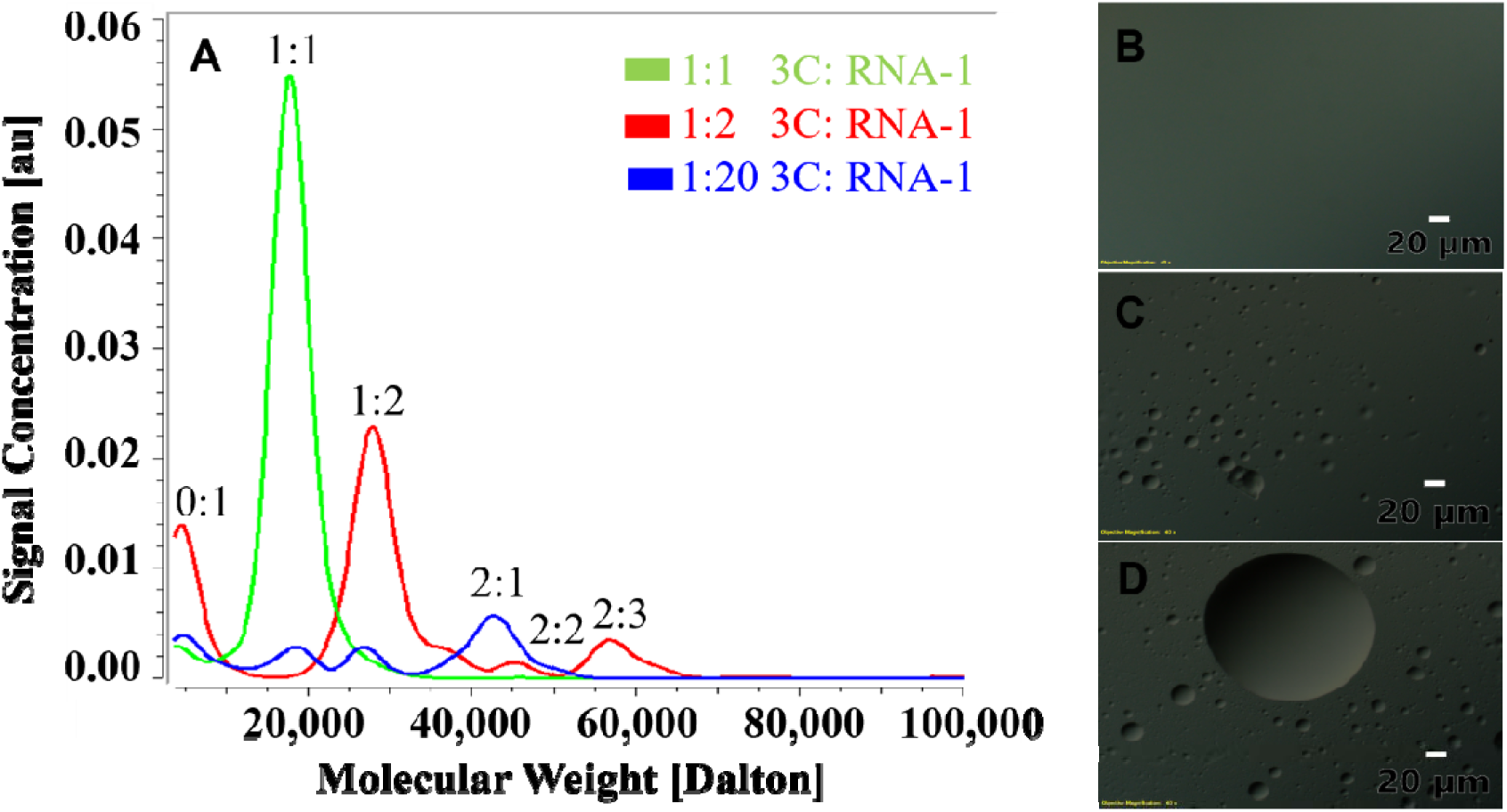
Interactions between PV-3C and RNA lead to multimeric complexes and liquid-liquid phase separation. (A) AUC of PV-3C with RNA-1, with PV-3C and RNA-1 forming a complex (green) at 1:1 molar stoichiometry. Upon increasing the RNA concentration (1:2 3C: RNA-1, red; 1:20 3C: RNA-1, blue), various multimolar-ratio complexes appear as indicated in red and blue, respectively. (B) Using DIC microscopy, LLPS is not apparent with 1:1 3C: RNA-1 complex. (C and D) However, at higher ratios (panel C, 1:2 3C: RNA-1; panel D, 1:20 3C: RNA-1 concentrations), DIC microscopy affirms the formation of the LLPS.

To confirm whether multimeric complex formation was a general phenomenon, we also tested RNA-2 (5’-CAUACUGUUGUAGGGGAA-3’ from oriR, **Table S1**) binding with 3C utilising NMR and AUC (**Fig. S18 and S19**). In contrast to the results with RNA-1, multimeric complexes involving 3C and RNA appear even at a 1:1 molar ratio (**Fig. S19**). This result is consistent with those from NMR that indicate the disappearance of amino acid peaks with the addition of RNA at lower RNA to protein ratios (**Fig. S18)**. At 1:0.5 molar ratio of 3C: RNA most of the peaks are absent from the SOFAST-HMQC spectra, which continues further to the complete absence of peaks at 1:1 molar ratio of 3C: RNA.

We also analyzed 3C and RNA binding using isothermal titration calorimetry (ITC)^25,26^. In this experimental set-up, PV-3C (at stock concentration of 400 µM) was titrated into RNA-1 (20 µM). The thermogram (**Fig. S20**) revealed two distinct binding events with different thermodynamic properties, consistent with two separate binding sites (Cluster-I and Cluster-II) from the NMR results (**Fig. 1**). The dissociation constants (∼52 nM and ∼950 nM) suggest substantially different binding affinities between two distinct binding sites. The two derived dissociation constants would also be consistent with the slow-exchange regime as observed by NMR (i.e. peak disappearance)^27,28^. Unfortunately, we were unable to collect ITC data with other RNA, likely due to precipitation challenges.

### Condensate formation observed from differential interference contrast microscopy

While AUC can detect changes in sedimentation behavior that suggest multi-molar ratio complex formation and/or phase separation, it does not provide direct visual confirmation. Differential interference contrast (DIC) microscopy has been widely employed to study liquid-liquid phase separation (LLPS)^29,30^. Here, DIC microscopy suggested the formation of microscopic phase-separated droplets owing to RNA addition to 3C, but only under conditions that lead to multimeric complex formation. For instance, the 1:1 3C: RNA-1 complex does not show evidence of phase separation (**Fig. 3B**), unlike those samples using 1:2 and 1:20 3C: RNA-1 ratios (**Fig. 3C and 3D**, respectively). Further addition of protein or RNA does not affect the process of LLPS. PV-3C by itself or RNA by itself does not lead to the same behavior; notably, 3C is predominantly in its monomeric form up to 50 µM concentration (**Fig. S17)**.

In contrast, even a 1:1 mixture of PV-3C and RNA-2 shows evidence of LLPS (**Fig. S19**), and several higher-order multimeric complexes are observed by AUC for this sample (**Fig. S19**). This finding suggests that RNA-2 is more capable of seeding oligomerization of PV-3C, leading to LLPS. In fact, LLPS was apparent for all other RNAs tested (**Fig. S21**).

### PV-3CD and RNA interactions also lead to LLPS

Having examined the effect of RNA on PV-3C, we also confirmed that addition of RNA-1 to PV-3CD likewise led to LLPS, as observed by DIC microscopy (**Fig. 4**), under all conditions tested, including at 1:0.5 and 1:1 mixture of RNA and protein. Similarly, all RNAs tested with 3CD led to LLPS according to DIC microscopy (**Fig. S22**). These findings were consistent with the observation of larger oligomers in the 20-40 nm range according to negative-stain electron microscopy (**Fig. S23**).

**Figure 4.**
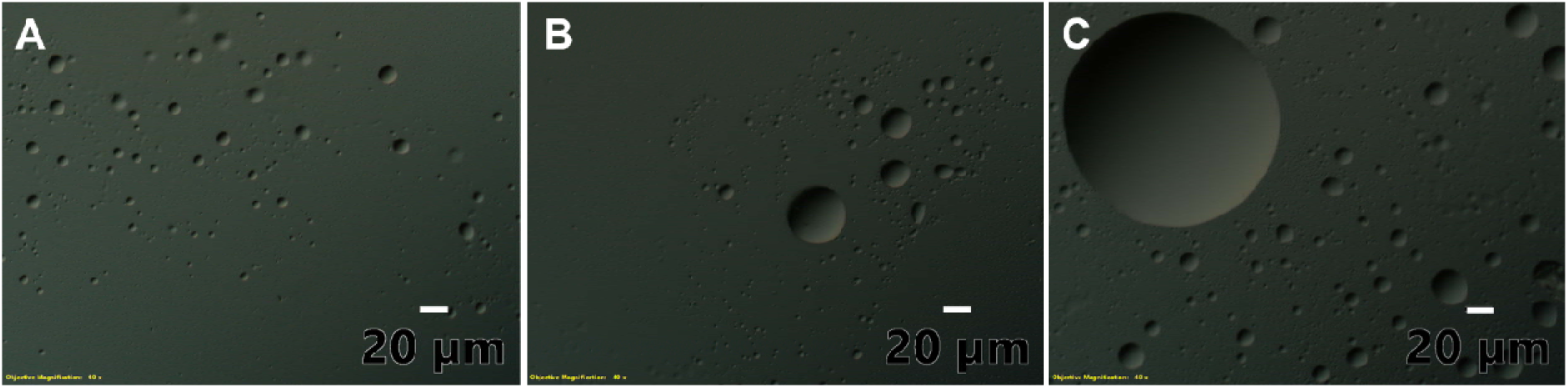
Interactions between PV-3CD and RNA lead to LLPS. DIC microscopy shows evidence for LLPS under all 3CD to RNA ratios tested (panel A, 1:0.5; panel B, 1:1; panel C, 1:2).

### Molecular model of the multivalent interactions between PV-3C/3CD and RNA

To gain a deeper insight into the multivalent nature of protein-RNA interactions, which are likely critical for LLPS, we performed molecular dynamics (MD) simulations of the PV-3C and PV-3CD proteins interacting with RNA-1 and RNA-2^31–33^. In our analysis, RNA was considered bound to the protein when any RNA heavy atom (non-hydrogen atom) was located within 5 Å of any protein residue. Using this geometric criterion, we defined a proximity probability (P_prox_) for each protein residue as P_prox_ = n/N, where n represents the number of timeframes during which any heavy atom of the given residue is bound to RNA, and N is the total number of frames in the MD trajectory. Thus, P_prox_ effectively quantifies the residence time of RNA molecules near protein binding sites throughout the simulation (**Figs. 5A, S24A)**.

**Fig 5.**
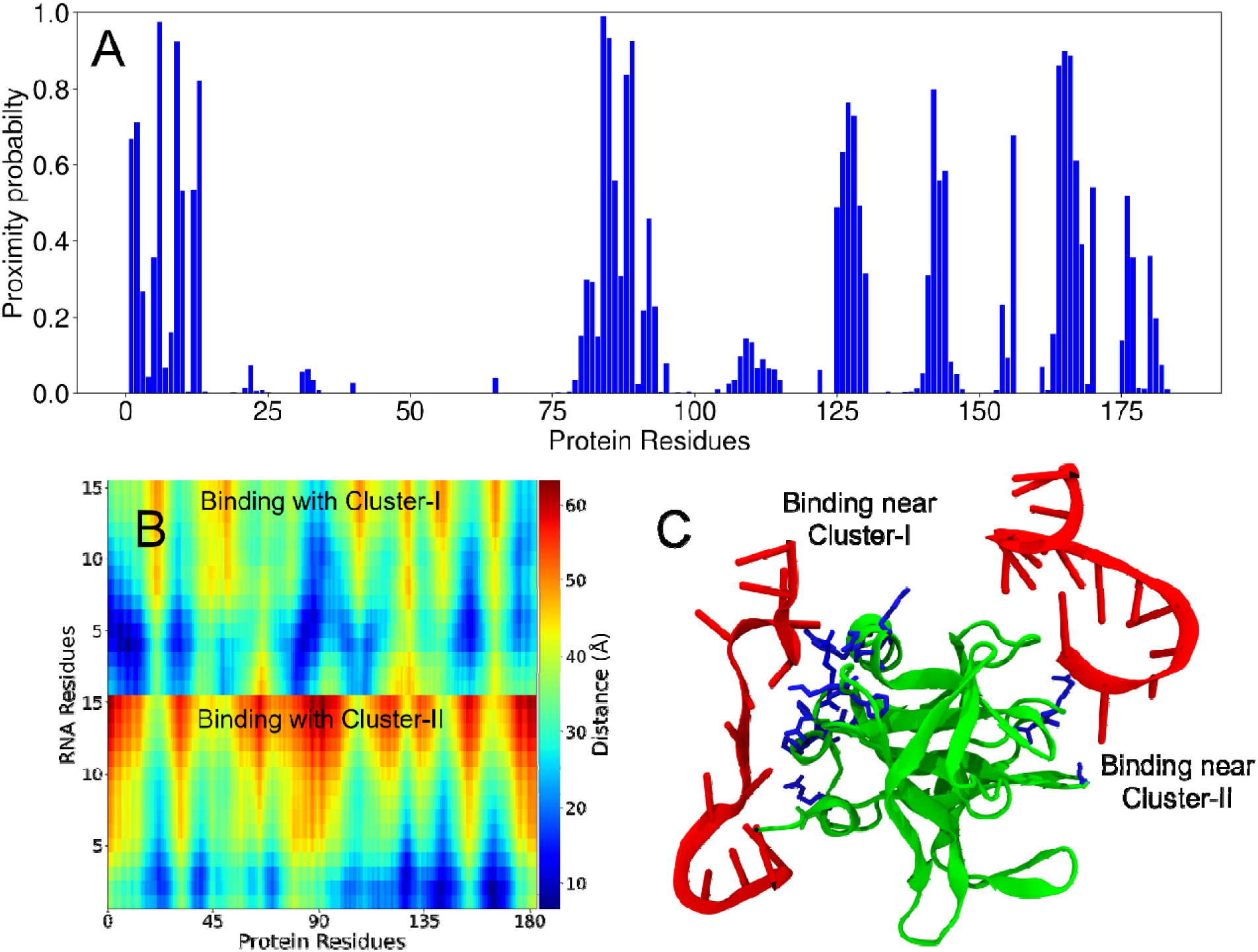
Two interaction sites between PV-3C protein and RNA-1 by MD simulations. (A) Proximity probabilities (*P_prox_*) of residues in 3C. (B) Distance map between the residues of RNA-1 and 3C. The color bar represents the color-coded distances underlying the distance map. The vertical axis shows the RNA residues of the two chains (each with 15 nucleotides). (C) Simulation snapshot showing two chains of RNA-1 interacting with the residues in Cluster-I and Cluster-II of the 3C protein (as also observed from NMR experiments). The protein and RNA chains are shown in green and red respectively. The residues colored blue have proximity probabilities greater than 0.5.

MD simulations encompass two primary types of non-bonded interactions: electrostatic (Coulomb potential) and van der Waals (Lennard-Jones potential). Protein-nucleotide interactions predominantly arise from electrostatic forces, driven by the negatively charged nucleotides interacting with positively charged protein regions (**Fig. 1C**)^34^. However, in this study, we exclusively used the geometric criterion described above to determine protein-RNA interactions and did not explicitly account for energetic contributions (because energy values are force field dependent). Consequently, a high P_prox_ value indicates protein regions frequently in proximity to RNA segments during the simulation. Such regions can include residues adjacent to the actual interaction sites, some of which may not exhibit significant chemical shift perturbations in NMR experiments. Therefore, direct one-to-one comparisons of interacting residues between computational and experimental results should be approached cautiously, as they may lead to misleading interpretations.

In the 3C simulations, both RNA-1 and RNA-2 exhibited substantial interactions with Cluster-I residues (**Fig. 5 and S24**). Both RNA oligonucleotides also interacted with Cluster-II residues, although the binding propensity for Cluster-II was apparently lower than that for Cluster-I (**Fig. 5A, S24A**). RNA-2 also displayed a higher binding propensity compared to RNA-1 for both residue clusters (**Fig. S24A**). Distance maps between RNA molecules and 3C also summarize the average interactions throughout the trajectory (**Fig. 5B, S24B**; blue indicating shorter distances between protein and the two RNA molecules). MD simulation videos provide a clear visual representation of the 3C and RNA interaction (**SV1 And SV2**).

Given that the MD simulations were largely consistent with the PV-3C NMR results, we extended this methodology to investigate RNA binding to full-length PV-3CD. While we have previously published NMR results with PV-3CD^4^, these studies were limited to Ile δ1-^13^CH_3_ resonances, which would provide limited information about RNA binding. Similar to the results with PV-3C, residues in Cluster-I and Cluster-II showed propensity to interact with RNA (**Fig. 6, S25**). Additionally, and perhaps not surprisingly given that 3D is the RNA-dependent RNA polymerase, RNA also interacted with the 3D domain of PV-3CD.

**Fig 6.**
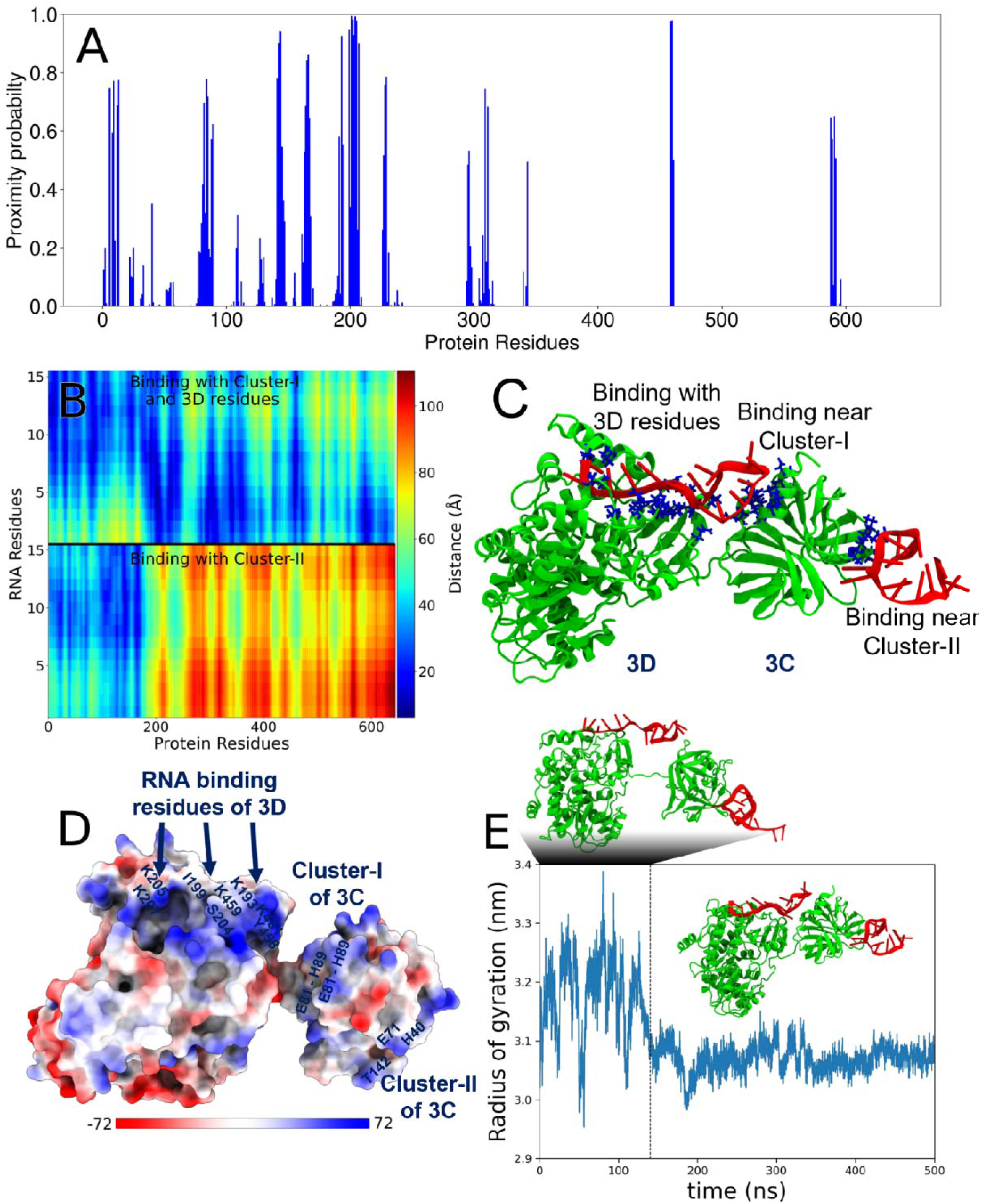
Interactions between PV-3CD protein and RNA-1 from MD simulations. (A) Proximity probabilities (*P_prox_*) of residues in 3CD. (B) Distance map between the residues of RNA-1 and 3CD. The color bar represents the color-coded distances underlying the distance map. The vertical axis shows the RNA residues of the two chains (each with 15 nucleotides). (C) Simulation snapshot showing two chains of RNA-1 interacting with the residues in Cluster-I and Cluster-II of the 3CD protein. The former also interacts with positively charged residues in the 3D domain. The protein and RNA chains are shown in green and red respectively. The residues colored blue have proximity probabilities greater than 0.5. (D) Electrostatic map of the PV-3CD protein surface. (E) Time evolution of the radius of gyration (Rg) of PV-3CD obtained from MD simulations. Representative snapshots of the protein structures in these two distinct states are presented along with the Rg time-trace.

Like 3C, the binding propensity for Cluster-II in 3CD was apparently lower than that for Cluster-I with both RNA oligonucleotides (**Fig. 6A, S25A**). Notably, there is a contiguous surface of positively-charged groups connecting 3C and 3D domains (**Fig. 6D**), likely helping to facilitate RNA binding due to the proximity of RNA-binding regions at the interface between these domains. Intriguingly, RNA binding also apparently induces proximity and compaction between the 3C and 3D units within the PV-3CD protein, as shown by a distinct structural transition, characterized by a significant decrease in Rg, is observed around 130 ns of the simulation (**Fig. 6E**). 3CD-RNA-1 forms a stable complex as concluded from Rg (**Fig. 6E**) because the stable complex exhibits a relatively constant Rg over the simulation time after binding, while fluctuations in Rg indicate dynamic regions or transient interactions within the complex. MD simulation videos provide a clear visual representation of the 3CD and RNA interaction (**SV3 and SV4**).

## DISCUSSION

Besides their proteolytic function, 3C and/or 3CD engage with RNA control elements in the picornavirus genome to help regulate and coordinate transcription and translation events^35,36^. Previous studies have brought insight into how PV-3C interacts with oriI, but little to no information has been available for similar interactions with oriL and oriR derived RNA^17^. It is especially noted that proposed RNA sites for 3C/3CD interaction have little to no sequence and/or structure similarities. As such, we investigated the structure and sequence specificity of RNA oligonucleotide binding to PV 3C and 3CD. While our studies revealed little specificity underlying 3C-RNA or 3CD-RNA interactions (at least using this set of RNA oligonucleotides), both sets of interactions can induce LLPS, a phenomenon that is important to various viral processes.

It is now well-established that LLPS is a complex process triggered by multivalent interactions, weak forces, and the interplay between proteins and RNA^31,32^. For 3C, NMR identified two potential regions of RNA interaction: Cluster-I including amino acid residues in the 80’s region (as identified in previous NMR studies) and the N-terminal alpha helix, and Cluster-II on the opposite face around/adjacent to the protease active site (**Fig. 1**)^15,37,38^. Interestingly, most of the identified residues belong to loops of PV-3C; the inherent flexibility in these loops may allow the residues to explore different conformations to allow for interactions with a variety of RNA sequences. Notably, both Cluster regions have positively charged surface potentials through which electrostatic interactions with negatively charged RNA could occur (**Fig. 1C**). Results with ITC (**Fig. S20**) and MD simulations (**Fig. 5**) are also consistent with two RNA binding sites, with Cluster-I likely interacting with a higher affinity than Cluster-II. ITC measurements revealed dissociation constants (K_d_) of approximately 52 nM and 950 nM (**Fig. S20**). Together with the MD simulations, these results might indicate that RNA-1 binds to Cluster-I with an affinity approximately 19-fold higher than to Cluster-II. At higher RNA concentrations, NMR peaks begin to disappear (**Fig. S16**), consistent with the formation of larger multimeric complexes, as verified by the SV-AUC results (**Fig. 3**). The connection between multivalency and LLPS formation is especially shown with the interactions involving 3C and RNA-1. With an equimolar mixture, there is only a 1:1 protein-RNA complex as observed by SAXS (**Fig. 2**) and SV-AUC (**Fig. 3A**). Under these conditions, there is no apparent LLPS by DIC microscopy (**Fig. 3B**). However, as RNA concentration is increased, multimeric complexes form (**Fig. 3A**), which likely lead to LLPS (**Fig. 3C-D**). In contrast, multimeric complexes using RNA-2 are apparent at the lower protein-RNA ratio (**Fig. S19**) and so is LLPS (**Fig. S19**). The computational results also suggest that RNA-2 has a higher propensity to interact with Cluster-I and especially with Cluster-II residues (**Fig. S24**). This phenomenon appears to be general as all other 3C-RNA interactions lead to decreases in NMR peak intensities (consistent with the formation of large multimeric complexes; **Fig. S2-S14**) and there is evidence for LLPS by DIC microscopy (**Fig. S21**). Moreover, this phenomenon likely extends to other picornaviral 3C proteins given their conservation of sequence (**Fig. S26)**, structure (**Fig. S27**) and electrostatic surface potential (**Fig. S28**). Altogether, our results suggest a mechanism by which interactions between RNA oligonucleotides and 3C induce LLPS (**Fig. 7**).

**Figure 7.**
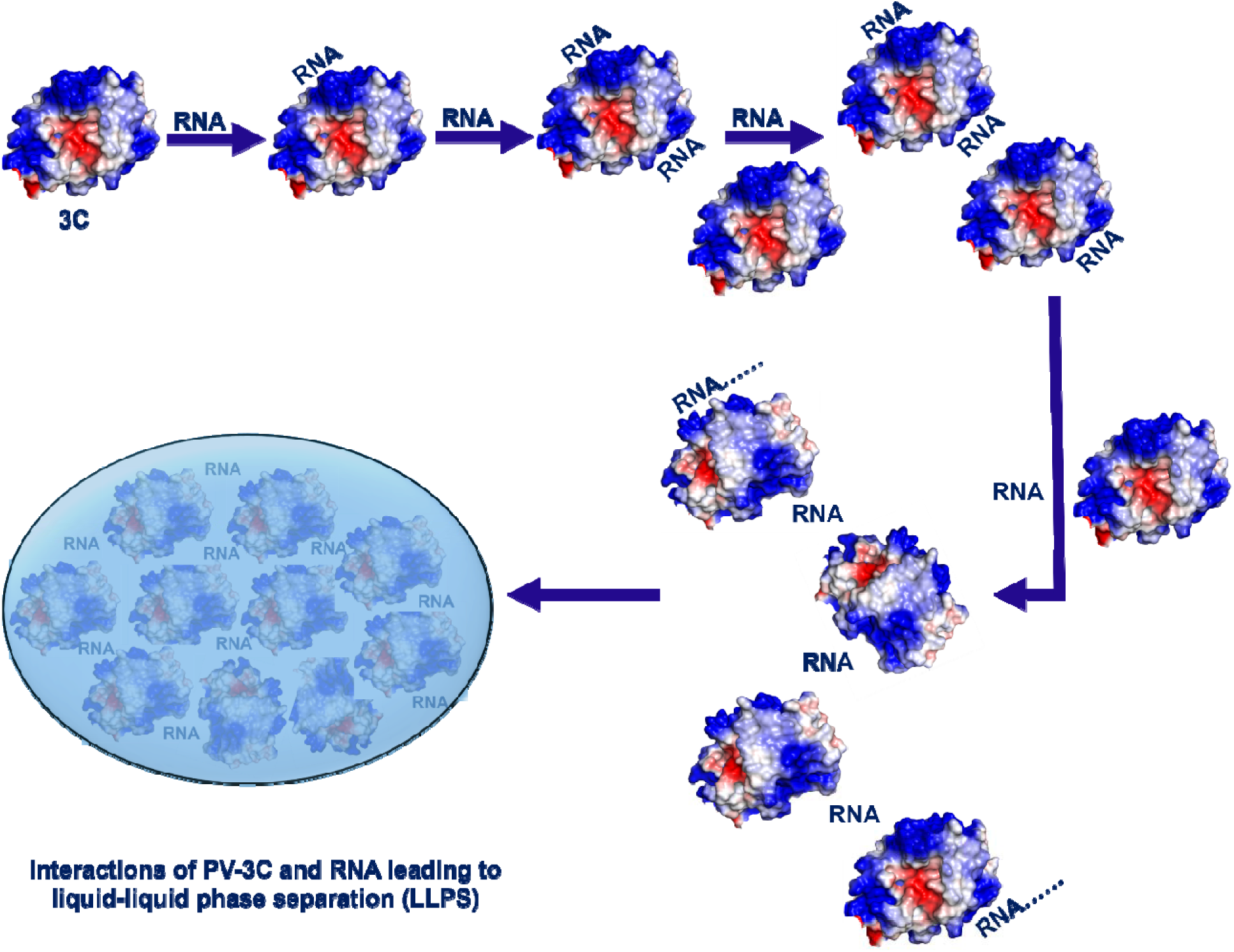
Multivalent interactions between 3C (or 3CD) and RNA oligonucleotides lead to liquid-liquid phase separation.

Given the results with 3C, it is then not surprising that RNA interactions with 3CD also lead to LLPS (**Fig. 4, S22**), as the 3D domain also interacts with RNA. MD simulations also suggest that RNA has propensity to interact with the corresponding Cluster-I and Cluster-II residues, and moreover, the Cluster-I RNA binding surface appears to be contiguous with an RNA binding surface on the 3D domain (**Fig. 6**). Given that 3CD is more abundant in cells than 3C, these results may be more biologically relevant^39^. In either case, protein-RNA-induced LLPS seems to be largely independent of the RNA sequence, such that 3C and/or 3CD could form such interactions with any RNA in the cell, not only those involving the CRE elements in the picornavirus RNA genome. These findings suggest that there may be mechanisms to prevent and/or leverage 3C/3CD-involved LLPS.

LLPS is increasingly recognized as a key component in driving the viral lifecycle, influencing viral replication, assembly, and host-pathogen interactions^40–42^. For example, LLPS may be involved in the formation of viral replication compartments, which may help in evading host cell immune surveillance and virus assembly, and interfere with host stress response^18,40–43^. For the latter, stress granules (SGs) and processing bodies (PBs) are both cytoplasmic ribonucleoprotein complexes that interact dynamically, exchanging mRNA and proteins to regulate translation, storage, and degradation of mRNA under stress conditions, including virus infection^44,45^. It is known that 3C expression inhibits SG formation and leads to PB dispersal, likely through its ability to cleave key protein factors G3BP1, important for nucleating SGs, and Dcp1a and PAN3, key components of PBs^46–48^. In contrast, the expression of PV-3CD has no apparent effect on SG assembly, although it has a similar effect on PB dispersal as PV-3C^47^. PV-3CD is also known to interact with many components of SGs, including eIF4G, GTPases, G3BP1, TIA1, PCBP and PABP^49–51^. It is also a key factor in the generation of virus replication organelles, although the relationship between these membrane-bound structures and the membraneless SGs and PBs is unclear^40,52,53^. As such, our results suggest further ways that 3C and/or 3CD might interact with protein-RNA condensates, potentially by first integrating into these complexes before cleaving essential proteins to cause disruption. LLPS, stimulated by interactions between RNA and 3C/3CD, could also promote the disassembly of SGs and PBs. Thus, the mechanisms revealed in this study hint towards possible functional and regulatory roles for 3C and/or 3CD in virally-induced LLPS-related events, including in stress responses.

## METHODS

### Materials

^15^N-ammonium chloride (NH_4_Cl) and D2O (D, 99.9%) were purchased from Cambridge Isotope Laboratories (Andover, MA, USA). All reagents were bought from VWR (Radnor, PA, USA) unless otherwise specified. The IS307 iSpacer” with a depth of 0.3 mm, was purchased from SunJin Lab Co. for use in Differential Interference Contrast (DIC) microscopy.

### Protein expression and isotopic labelling

pSUMO plasmids encoding C-terminal hexa-histidine tagged PV 3C, 3D, and 3CD were transformed into *Escherichia coli* BL21(DE3) pRARE cells. In 3D and the 3D domain of 3CD, the following amino acid substitutions were present to avoid aggregation ((compared to type 1 Mahoney strain): L446D and R455D^4,15,37^. On 3C and in the 3C domain of the 3CD, the following amino acid substitutions were included to inhibit aggregation and disrupt the intermolecular cleavage of 3CD: E55A, D58A, E63A and C147A. The aggregation-preventing mutations hinder oligomerization interfaces on 3C and 3D, enabling large concentrations of 3CD required for NMR and other biophysical studies.

Following transformation, cell cultures were shaken in 10 mL M9 minimal media (6.0 g/L Na_2_HPO_4_, 3.0 g/L KH_2_PO_4_, 0.5 g/L NaCl, 2 g/L glucose, 1.0 g/L NH_4_Cl, 0.1 mM CaCl_2_, 1 mM MgSO_4_, trace metal mix (Teknova, Hollister, CA, USA), MEM vitamin mix (Thermo Fisher, Bellefonte, PA, USA), 50 g/mL kanamycin, and 30 g/mL chloramphenicol) for 16–20 h until the optical density at 600 nm (OD600) was higher than 0.6. For ^15^N labelling, 1 mL of the 10 mL culture was inoculated to 50 mL of M9 medium and shaken overnight at 200 rpm at 30°C. The next morning, 20 mL of the overnight culture was used to inoculate 1 L of M9 media (including 1 g/L ^15^N NH_4_Cl from Cambridge Isotope Laboratories, 2 mM MgSO_4_, 12.0 g/L Na_2_HPO_4_, 6.0 g/L KH_2_PO_4_). Protein expression was initiated with 1 mM isopropyl--D-1-thiogalactopyranoside (IPTG) once OD600 reached 0.6–0.8, and the cultures were then shaken at 25_°_C. After 16-20 hours, cells were harvested through centrifugation (3900 xg, 30 min, 4 C), rinsed with a buffer (10 mM Tris, 1 mM EDTA pH 8) and again centrifuged (3800 xg, 15 min, 4 C). Solutions were then decanted and the resultant pellets weighed before storing at -80°C. Unless otherwise mentioned, all the cultures were incubated at 37 °C while shaking at 200–250 rpm.

### Purification of PV-3C and PV-3CD

Cell pellets with expressed PV-3C were resuspended in 20 mM HEPES, 50 mM NaCl, 1mM EDTA, 5mM β-mercaptoethanol, 5 mM imidazole, pH 7.5, with 1.4 g/mL pepstatin A, 1 g/mL leupeptin. Cell pellets with expressed PV-3CD were resuspended in a buffer containing 100 mM potassium phosphate, 10 mM β-mercaptoethanol, 120 µM ZnCl_2_, 20% glycerol, pH 8.0, with 1.4 g/mL pepstatin A, 1 g/mL leupeptin, 500 mM phenylmethanesulfonyl fluoride (PMSF). The cells were then lysed, followed by polyethyleneimine (PEI) precipitation and ammonium sulfate precipitation to 60% saturation. The ammonium sulfate pellet was resuspended, followed by the purification of proteins by nickel-nitrilotriacetic acid (Ni-NTA) affinity chromatography, following protocols outlined in refs^15,37,54^. Following the cleavage of the SUMO tag by 1–2 g ubiquitin-like specific-protease (ULP-1), samples were dialyzed against 100 mM potassium phosphate (pH 8.0), 100 mM sodium chloride, 60 mM ZnCl_2_, 5 mM β-mercaptoethanol and 20% glycerol. The samples were then concentrated using 30 kD molecular weight cutoff (3CD) or 10 kD molecular weight cutoff (3C) Sartorius Vivaspin spin concentrators in a buffer containing 10 mM HEPES, 50 mM NaCl, pH 7.50.

### NMR sample preparation

To prepare ^15^N labelled protein samples, we utilised the buffer containing 10 mM HEPES pH 7.5 and 50 mM NaCl with 10% D_2_O for deuterium lock. The protein samples were concentrated using a 0.5 mL Millipore Amicon Ultra Centrifugal Filters, 3K MWCO. The protein concentration was calculated by measuring the absorbance at 280 nm (The molar extinction coefficient (ε) for 3C =8960 M^−1^ cm^−1^, 3CD = 84690 M^−1^ cm^−1^). All NMR experiments were carried out on a 600 MHz Bruker NEO spectrometer with 5mm TCI single-axis gradient cryoprobes (Bruker, Billerica, MA, USA). NMR data was processed using Sparky software^55^. The ^1^H-^15^N SOFAST-heteronuclear multiple quantum coherence (HMQC) spectra were run with 32 ns, 128 points were used in the indirect dimension (F1) and 2048 points were used in the direct dimension (F2)^21,56–59^.

### Sedimentation velocity analytical ultracentrifugation (AUC)

PV-3C and PV-3CD, both with and without RNA-1 and/or RNA-2, were loaded into 3 mm epon-charcoal centerpieces sandwiched between sapphire windows for analytical ultracentrifugation analysis^60^. The cells were loaded into either an An50 or An60 titanium rotor (depending on number of samples for the run) and the rotor was then placed in the vacuum chamber of a Beckman-Coulter Optima multiwavelength analytical ultracentrifuge. The vacuum chamber was evacuated and then the rotor and samples were allowed to equilibrate to the experimental temperature of 25°C for ∼ 2 hours. Once equilibrated, a method was written for the run in the UltraScan III software^61^. For this method, the rotor accelerated to 42,000 RPM, and radial scans of each sector were performed every 2 minutes for 16 hours. For AUC measurements, the wavelength used for 3C was 280 nm, and 290 nm for 3CD. 280 nm is the most used wavelength for protein detection in AUC due to the presence of aromatic amino acids. Therefore, we chose a wavelength of 280 nm for the AUC measurement of 3C (MW=20.69 kDa), whereas the signal was too high for 3CD at this wavelength because 3CD (MW=71.92 kDa) is a larger protein, resulting in higher absorption. As a result, we picked 290 nm for the AUC measurement of 3CD to keep the absorbance at or below 1. Once the run was completed, the data were analyzed using UltraScan III^61^. The data were first converted from raw radial intensity data to pseudo-absorbance data. The scans were then fit to solutions to the Lamm equation, with additional fitting to account for time-independent noise. Once the RMSD for the fits were <0.003, the data were re-fit to determine the correct meniscus position for each sample and to account for radially invariant noise as well. Then, a final, iterative fit was performed. The final S-value range was 1-10 with a resolution of 100, and the final frictional ratio range was 1-4 with a resolution of 64.

### Small angle X-ray scattering (SAXS)

SAXS experiments were conducted in a buffer comprising 10 mM HEPES pH 7.5 and 50 mM NaCl. BioSAXS data were collected using X-rays generated by a Rigaku MM007 rotating anode X-ray source, in conjunction with the BioSAXS2000^nano^ Kratky camera system^61^. This system is equipped with OptiSAXS confocal max-flux optics, specifically designed for SAXS, and features a highly sensitive HyPix-3000 Hybrid Photon Counting detector.

The sample-to-detector distance was set at 495.5 mm and calibrated using silver behenate powder from ‘The Gem Dugout’ company (State College, PA). The usable momentum transfer scattering vector range, spanned from q_min_= 0.008 Å ¹ to q_max_= 0.6 Å ¹ (with q = 4πsin(θ)/λ, and 2θ representing the scattering angle). The X-ray beam wavelength was 1.54 Å, with a Kratky block attenuation of 22%, and a beam diameter of approximately 100 μm.

Samples were introduced using the Rigaku autosampler into a quartz capillary flow cell situated on a sample stage cooled to 4°C^61^. The entire X-ray flight path, including the beam stop, was maintained under vacuum < 1×10 ³ torr to minimize air scatter. Automated data collection, incorporating thorough cleaning cycles between samples, was performed using the Rigaku SAXSLAB software. Data sets were collected for 60 minutes with six ten-minute images using the autosampler quartz flow cell. Buffer SAXS data collected over 60 minutes using the same flow cell and was used for the reference subtraction. The overlays of the six ten-minute images that were averaged in each case showed that there was no X-ray radiation damage. Three replicate SAXS data were seen to overlay well and were further averaged to get the final SAXS curve. The ATSAS Primus software was employed to determine the forward scattering I(0) and the radius of gyration (Rg) using the Guinier approximation. The Guinier approximation assumes that at very small angles (q <1.3/Rg), the intensity follows the formula I(q) = I(0)exp[−1/3(qRg)²]. These size parameters were consistent with measurements obtained from the 3C crystal structure model. The data files underwent analysis for the radius of gyration, Dmax, Guinier fits, Kratky plots, and pair distance distribution function^62^.

High-q Kratky plots indicated well-folded proteins with no flexibility for the 3C protein and some for the RNA complex. The pair-distance distribution function P(r) was calculated using GNOM, yielding the maximum particle dimension (Dmax)^62^. Solvent envelopes were generated from DENSS, an algorithm used for calculating ab initio electron density maps directly from solution scattering data. The solvent envelope for the 3C protein alone SAXS mostly matched the monomer model. The RNA and 3C model was built guided by the residues identified by the NMR chemical shifts. The 3C-RNA complex had an envelope agreeing with a 1:1 3C: RNA complex stoichiometry and was missing the flexible RNA termini. Theoretical scattering profiles of the 3C crystal structure model and the constructed model for the complex with RNA were computed and fitted to experimental scattering data using CRYSOL, resulting in good Chi-square fits.

### Differential interference contrast (DIC) microscopy

DIC microscopy was performed utilizing an Olympus BX 61 microscope at the Microscopy Core Facility (Huck), Pennsylvania State University. We utilized the biological configuration with automated 4-color plus DIC image collection, polarization, darkfield, and brightfield^29^. The illumination happens through Mercury vapour, provided through either a rapid automated shutter or conventional filter cubes. The objective of the Biological Configuration includes UplanFL 40X/0.75 for our images. All objectives in this microscope have DIC optics. The Digital cameras are Hamamatsu cooled digital cameras (ORCA ER, Model C4742-80) and Olympus DP71. The prior controller with the joystick controls focus and location. CellSens software was used for image processing. PV-3C and PV-3CD samples with RNA in the HEPES buffer (10 mM HEPES pH 7.5 and 50 mM NaCl) were prepared, and a drop was mounted on clean glass slides with a coverslip for high-quality DIC microscopy imaging. There was no staining, and appropriate control of buffer, protein, and RNA were recorded before observing LLPS in the protein-RNA samples.

### Isothermal Titration Calorimetry

ITC data were collected using a TA Affinity ITC automated microcalorimetry instrument with a stirring rate of 125 rpm at 25°C. For each experiment, 25 injections of 2.0μL volume were performed^25^. Concentrations were 400µM of 3C into 20µM RNA. A second titration set was performed: 400µM of 3C into the buffer (10 mM HEPES pH 7.5 and 50 mM NaCl) for baseline subtraction. Data analyses were performed using the NanoAnalyze software version (4.0.2). Data were fit using a two-site binding model.

### Negative staining electron microscopy

For visualization by transmission electron microscopy (TEM), the samples were subjected to negative staining. A 3.5 μL aliquot of the sample was applied to a carbon support TEM grid (400 mesh, Ted Pella) and incubated for 1 minute^63^. Subsequently, excess liquid was carefully blotted away using filter paper. The grid was then washed twice with 10 μL of ultrapure water to remove any unbound material. The grid was then stained with 10 μL of freshly prepared uranyl formate solution (0.7% w/v, pH 4.5) for 30 seconds. Finally, after blotting away excess staining solution, the grid was air-dried at room temperature.

The images were recorded using FEI Titan Krios at the Huck Institute of the Life Sciences at Penn State. We would like to thank Dr Sung Hyn Cho for helping us with the experiment.

### Molecular dynamics (MD) simulation: system preparation

The initial coordinates of the PV-3C and PV-3CD proteins were obtained from the Protein Data Bank (PDB: 1L1N and 2IJD, respectively)^6,11,64^. The last three residues (Gln181, Ser182, and Gln183) were missing from the crystallographic PDB structure of 3C. As such, these were modelled using Pymol-2.5.0 and refined using the ModLoop webserver of Modeller^65,66^. The initial structures of the RNA fragments of RNA-1 (GGC GGC GUA CUC CGG) and RNA-2 (CAU ACU GUU GUA GGG GAA) were generated using the NAflex web server^67^. Two chains of each RNA were put in a cubic box of sides ∼11 nm and ∼15 nm, along with the PV-3C and PV-3CD, respectively, resulting in a total of four protein-RNA systems. The cluster-I region (see Fig. 1) readily interacts with the RNAs, whereas the cluster-II region (see Fig. 1) shows lower RNA-binding propensity. Hence one of the RNA chains was randomly placed in the simulation box, whereas the second chain was placed in the vicinity of cluster-II.Each system was solvated with a 3-point water model. Approximately 50000 and 100000 water molecules were added to the systems containing PV-3C and PV-3CD, respectively. Sodium ions were added to neutralize the systems. Salt at 50 mM NaCl was added to mimic experimental conditions. The protein, RNA and ions were modelled using the CHARMM36m force field and the CHARMM-modified TIP3P model was used to describe the water molecules^68^.

### MD simulation runs and analysis

All MD simulations were performed using the GROMACS-2022.4 simulation package^69^. The following simulation protocol was applied to both PV-3C and PV-3CD systems. An initial energy minimization was performed using the steepest descent algorithm to get rid of unrealistic atomic contacts. This was followed by a short 1 ns equilibration under NpT conditions (temperature (T) = 300 K and pressure (p) = 1 bar) with harmonic position restraints on the heavy atoms of the protein and the RNA (with a force constant of 1000 kJ mol^−1^ nm^−2^) to let the water and ions equilibrate. Thereafter, the position restraints were lifted, and the system was allowed to equilibrate under NVT conditions (T = 300 K) for 10 ns. This was followed by a 500 ns production run under similar NVT conditions, with the data being saved at a frequency of 10 ps.

All atomistic MD simulations were performed using the leap-frog integrator with a time step of 2 fs. The v-rescale (stochastic velocity rescaling) thermostat and the c-rescale (stochastic exponential relaxation) barostat were used to control temperature and pressure (wherever necessary), respectively^70^. Protein bonds involving hydrogen atoms and the internal degrees of water molecules were constrained using the LINCS and SETTLE algorithms, respectively^71^. Short-range electrostatic and Lennard-Jones interactions were calculated up to a distance of 1.0 nm. Long-range electrostatic interactions were treated with the particle-mesh Ewald technique with a grid spacing of 0.12 nm^72^.

## Acknowledgements

The study was supported by NIH (R01AI104878 to DDB; S10-OD028589 for SAXS and S10-OD032215-01 for the Optima AUC to NY). We would like to thank Boehr lab members Alyson Boehr, Jie Yu, Mohammad Hasan and visiting faculty Dr. Sara Khan (CUI Abbottabad) for their technical assistance and project feedback, Julia Fecko (Biomolecular Interactions Facility at Penn State) for assistance with ITC, John Catolina (Microscopy Facility at Penn State) for assistance with DIC, Drs. Sung Hyun Cho and Ibrahim Moustafa (Cryo-electron Microscopy Facility at Penn State) for assistance with EM, and Drs. Tapas Mal and Christy George (NMR Facility in the Department of Chemistry, Penn State University) for assistance with NMR experiments. We also thank the Max Planck Computing and Data Facility and Prof. Gerhard Hummer for computational support.

