## Supplemental Information for "Interactions between the picornavirus 3C(D) main protease and RNA induce liquid-liquid phase separation"

**Table S1.** List of RNA sequences utilised for experiments with the origin

| **RNA name** | **Sequence** | **Origin** |
| --- | --- | --- |
| RNA-1 | GGCGGCGUACUCCGG | oriL- of PV-genome |
| RNA-2 | CAUACUGUUGUAGGGGAA | oriR- of PV-genome |
| RNA-3 | UUUUUUUUUUUUUUU |  |
| RNA-4 | ACCCCAGAGGCCCAC | oriL- of PV-genome |
| RNA-5 | CCAUCAAGAAUCCUA | SARS-COV2 genome |
| RNA-6 | AAAAAAAAAAAAAAA |  |
| RNA-7 | CGCAGCAAACACAAG | PV-RNA |
| RNA-8 | CUACCUCAGUCGAAU | oriR- of PV-genome |
| RNA-9 | CGUUGGCUUGACUCA | oriR- of PV-genome |
| RNA-10 | GGGGUUGUACCCACCCC | oriL- of PV-genome |
| RNA-11 | CUCCGGUAUUGCGGUACCCUUG | oriL- of PV-genome |
| RNA-12 | GAUUGGGUCAUACUGUUGUAG | oriR- of PV-genome |
| RNA-13 | CAGAGGAACACGUGGCGGCG | oriL- of PV-genome |
| RNA-14 | UUUUCUUUAAUUCGGAGAAAA | oriR- of PV-genome |
| RNA-15 | AGUUCAAGAGC | PV-genome |

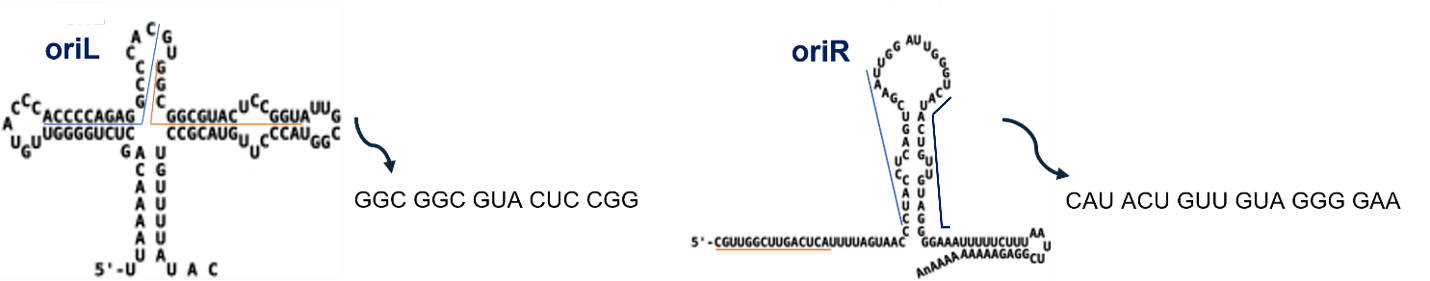

**Figure S1.** **Predicted secondary structure of oriL and oriR from the PV RNA genome.** RNA-1 and RNA-2 were based in part on RNA sequences from oriL and oriR respectively.

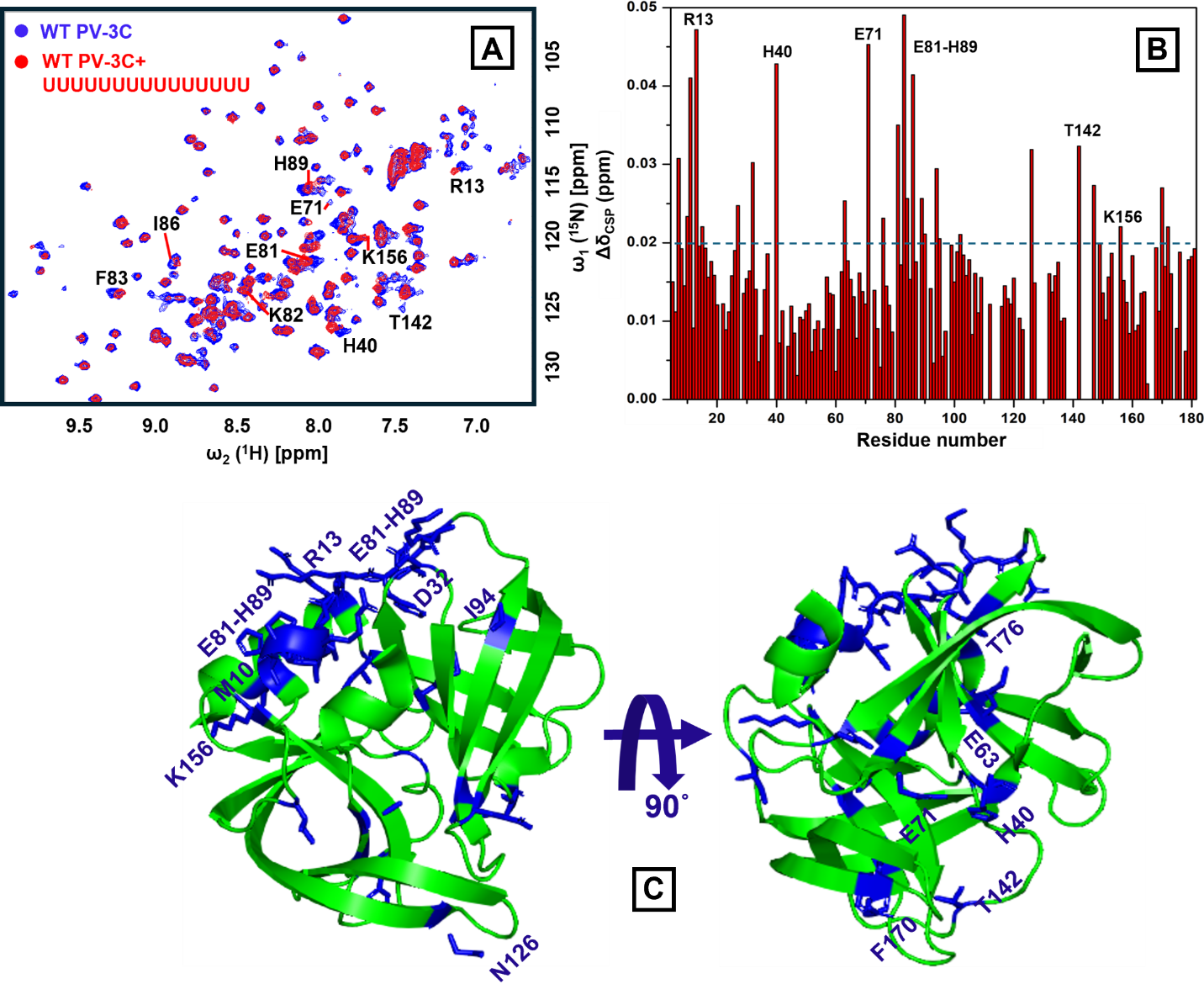

**Figure S2. PV-3C and RNA-3 (UUUUUUUUUUUUUUU) binding.** (A) ^1^H-^15^N SOFAST-HMQC spectra were compared with free wild-type PV-3C (blue) and RNA-3 bound PV-3C (red) in 1:0.5 stoichiometry; the residues with substantial chemical shift perturbation (CSP) are labelled in the spectra. (B) NMR CSP values for the amino acid residues of PV-3C are plotted; NMR CSPs were calculated using the following equation Δδ_combined_ = (Δδ_H_^2^ + (Δδ_N_/5)^2^)^0.5^, where Δδ_H_ and Δδ_N_ are the chemical shift differences between 3C with and without RNA for the backbone amide proton and nitrogen respectively. (C) Residues showing substantial CSPs in the presence of RNA are represented in blue on the 3C X-ray crystal structure (PDB ID: 1L1N) using PyMOL. These residues include cluster-I and cluster-II residues as determined using RNA-1 (see Figure 1). In addition, M27 and D32, near cluster-I, and E63, N126, and F170, near cluster-II, also show CSP. The PV-3C protein concentration was 25 µM and RNA was added following a 1:0.5 stoichiometry. Both protein and RNA were in a buffer containing 10 mM HEPES, 50 mM NaCl, and pH 7.5. The ^1^H-^15^N SOFAST-HMQC NMR experiment was recorded in a 600 MHz Bruker NEO spectrometer equipped with z-gradient triple resonance (^1^H, ^13^C, ^15^N) TCI cryogenic probe at 25˚C.

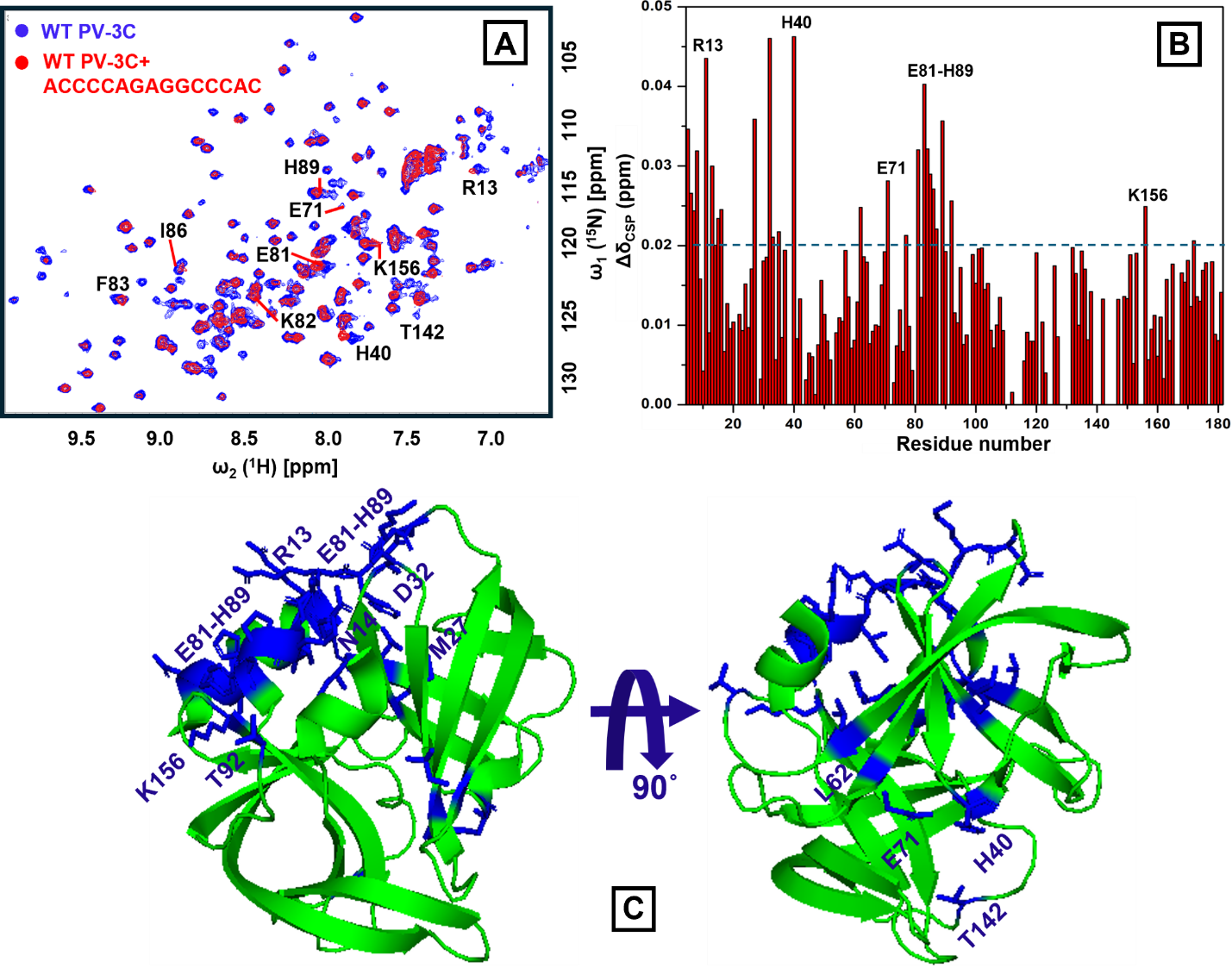

**Figure S3. PV-3C and RNA-4 (ACCCCAGAGGCCCAC) binding.** (A) ^1^H-^15^N SOFAST-HMQC spectra were compared with free wild-type PV-3C (blue) and RNA-4 bound PV-3C (red) in 1:0.5 stoichiometry; the residues with substantial CSP are labelled in the spectra. (B) NMR CSP values for the amino acid residues of PV-3C are plotted, using the equation in Fig. S2. (C) Residues showing substantial CSPs in the presence of RNA are represented in blue in the 3C X-ray crystal structure (PDB ID: 1L1N) using PyMOL These residues include cluster-I and cluster-II residues as determined using RNA-1 (see Figure 1). In addition, N14, M27, D32 and T92, near cluster-I, and L62, near cluster-II, also show CSP. Sample and NMR spectrometer conditions were as in Fig. S2.

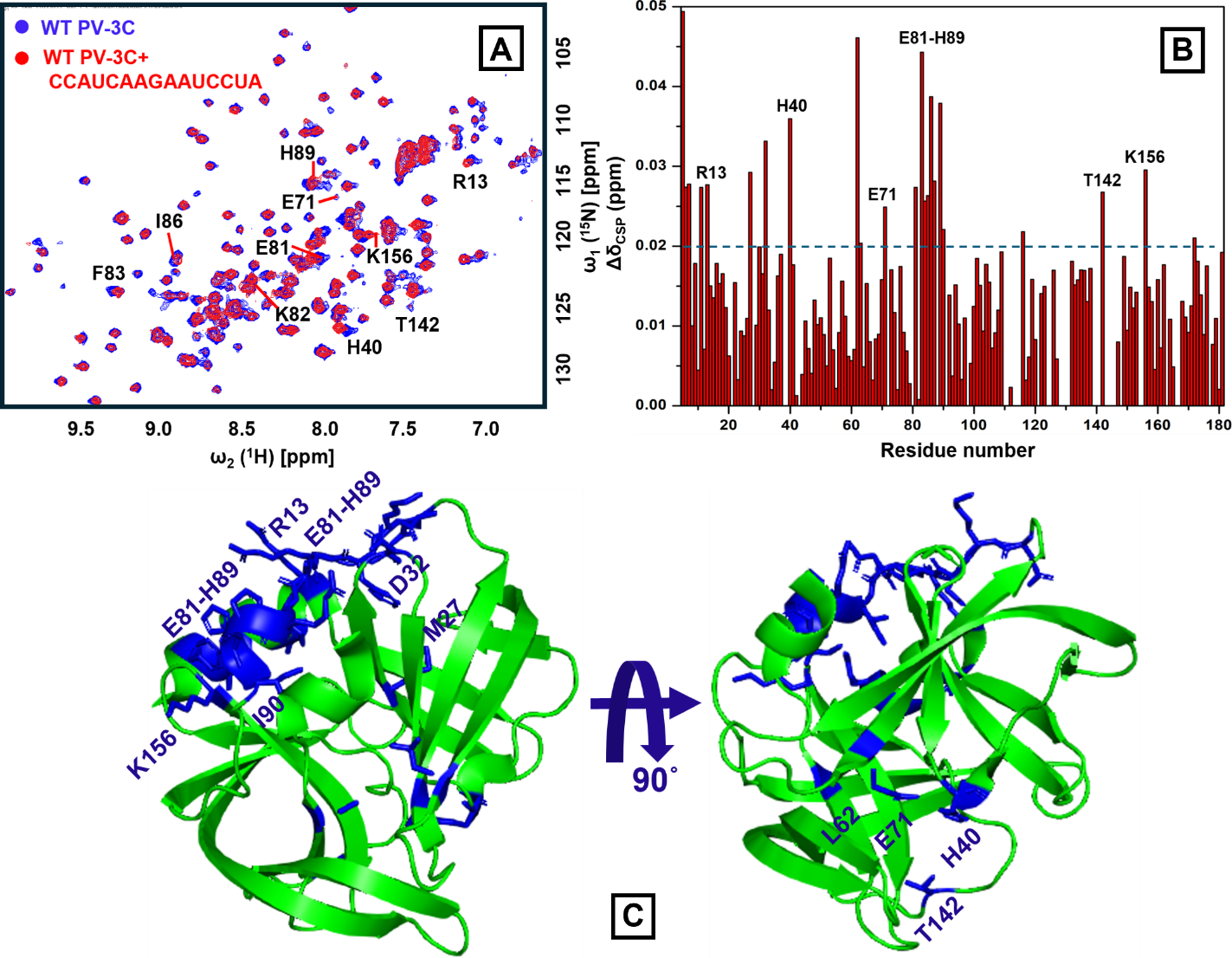

**Figure S4. PV-3C and RNA-5 (CCAUCAAGAAUCCUA) binding.**  (A) ^1^H-^15^N SOFAST-HMQC spectra were compared with free wild-type PV-3C (blue) and RNA-5 bound PV-3C (red) in 1:0.5 stoichiometry; the residues with substantial CSP are labelled in the spectra. (B) NMR CSP values for the amino acid residues of PV-3C are plotted, using the equation in Fig. S2. (C) Residues showing substantial CSPs in the presence of RNA are represented in blue in the 3C X-ray crystal structure (PDB ID: 1L1N) using PyMOL These residues include cluster-I and cluster-II residues as determined using RNA-1 (see Figure 1). In addition, M27, D32 and I90 near cluster-I, and L62, near cluster-II, also show CSP. Sample and NMR spectrometer conditions were as in Fig. S2.

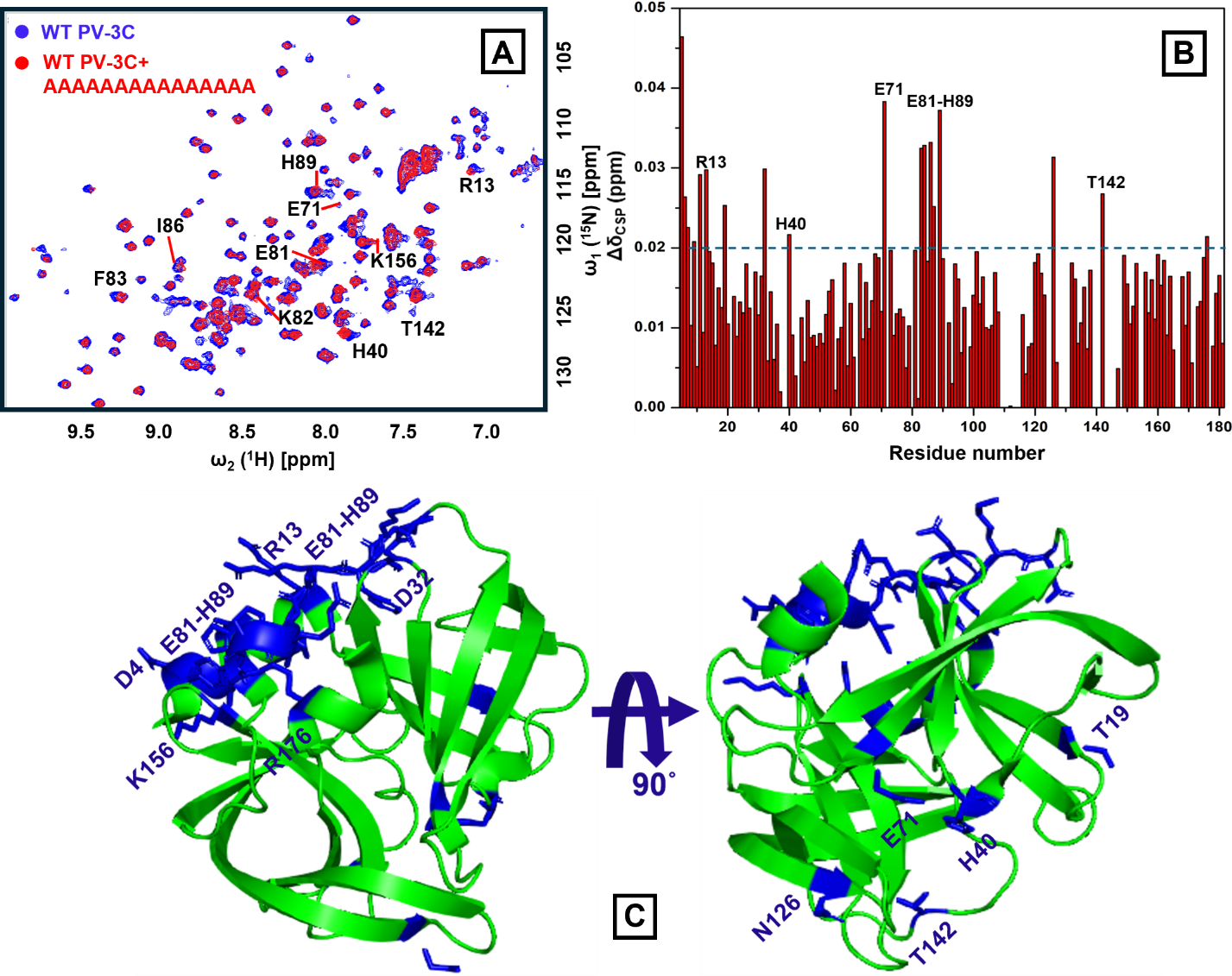

**Figure S5. PV-3C and RNA-6 (AAAAAAAAAAAAAAA) binding.** (A) ^1^H-^15^N SOFAST-HMQC spectra were compared with free wild-type PV-3C (blue) and RNA-6 bound PV-3C (red) in 1:0.5 stoichiometry; the residues with substantial CSP are labelled in the spectra. (B) NMR CSP values for the amino acid residues of PV-3C are plotted, using the equation in Fig. S2. (C) Residues showing substantial CSPs in the presence of RNA are represented in blue in the 3C X-ray crystal structure (PDB ID: 1L1N) using PyMOL These residues include cluster-I and cluster-II residues as determined using RNA-1 (see Figure 1). In addition, D32 and R176, near cluster-I, and T19 and N126, near cluster-II, also show CSP. Sample and NMR spectrometer conditions were as in Fig. S2.

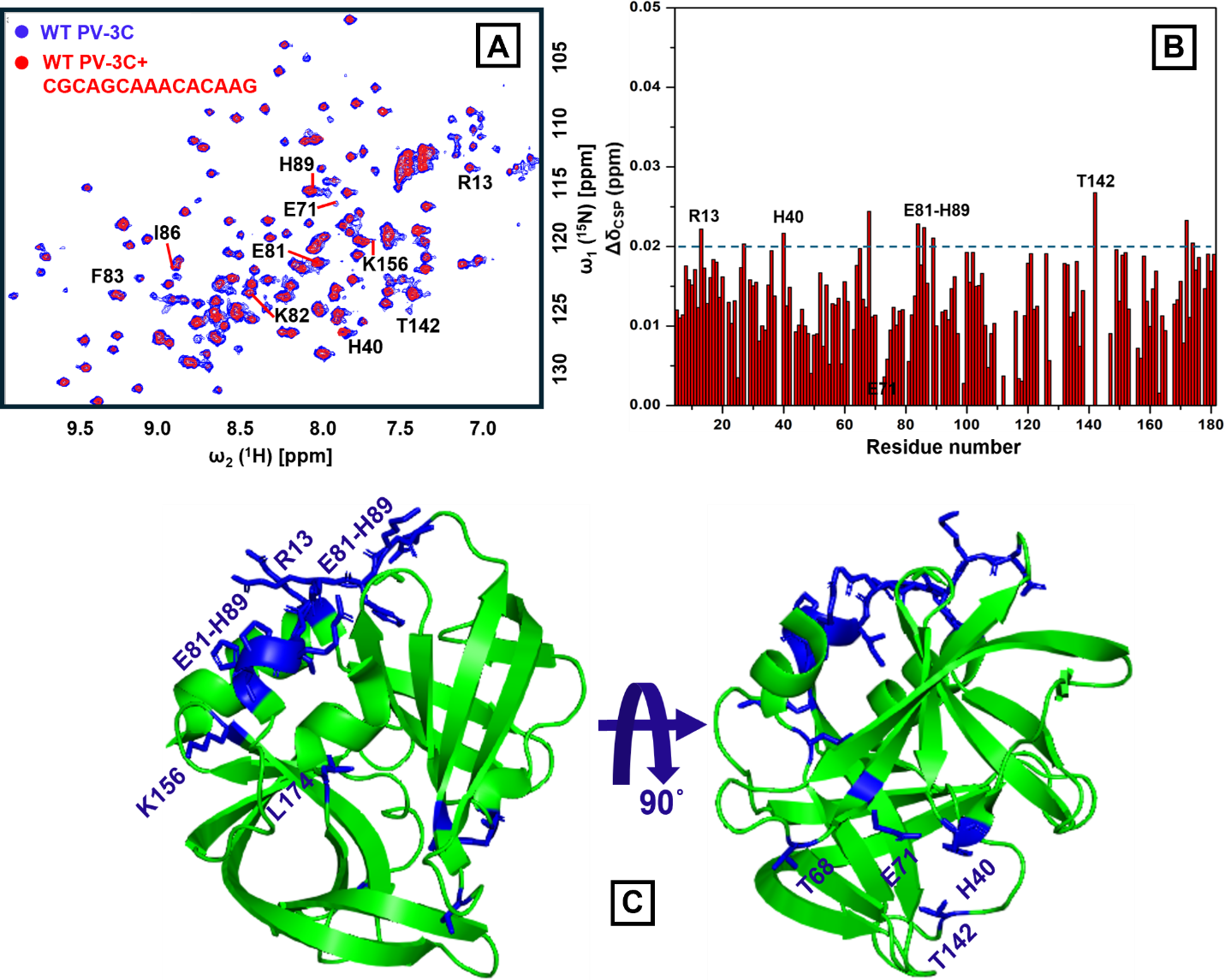

**Figure S6. PV-3C and RNA-7 (CGCAGCAAACACAAG) binding.** (A) ^1^H-^15^N SOFAST-HMQC spectra were compared with free wild-type PV-3C (blue) and RNA-7 bound PV-3C (red) in 1:0.5 stoichiometry; the residues with substantial CSP are labelled in the spectra. (B) NMR CSP values for the amino acid residues of PV-3C are plotted, using the equation in Fig. S2. (C) Residues showing substantial CSPs in the presence of RNA are represented in blue in the 3C X-ray crystal structure (PDB ID: 1L1N) using PyMOL These residues include cluster-I and cluster-II residues as determined using RNA-1 (see Figure 1). In addition, L174, near cluster-I, and T68, near cluster-II, also show CSP. Sample and NMR spectrometer conditions were as in Fig. S2.

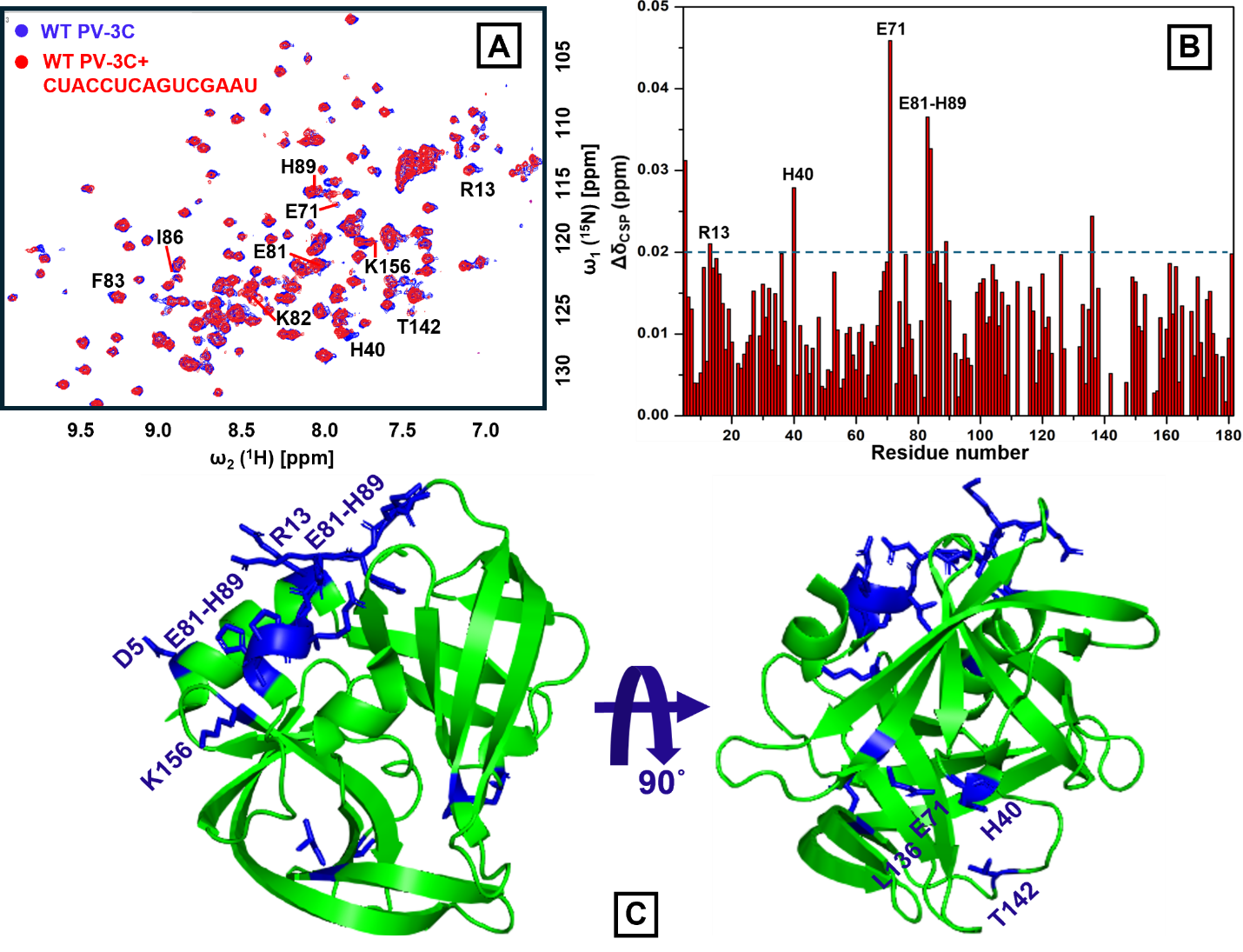

**Figure S7. PV-3C and RNA-8 (CUACCUCAGUCGAAU) binding.** (A) ^1^H-^15^N SOFAST-HMQC spectra were compared with free wild-type PV-3C (blue) and RNA-8 bound PV-3C (red) in 1:0.5 stoichiometry; the residues with substantial CSP are labelled in the spectra. (B) NMR CSP values for the amino acid residues of PV-3C are plotted, using the equation in Fig. S2. (C) Residues showing substantial CSPs in the presence of RNA are represented in blue in the 3C X-ray crystal structure (PDB ID: 1L1N) using PyMOL These residues include cluster-I and cluster-II residues as determined using RNA-1 (see Figure 1). In addition, L136, near cluster-II, also shows CSP. Sample and NMR spectrometer conditions were as in Fig. S2.

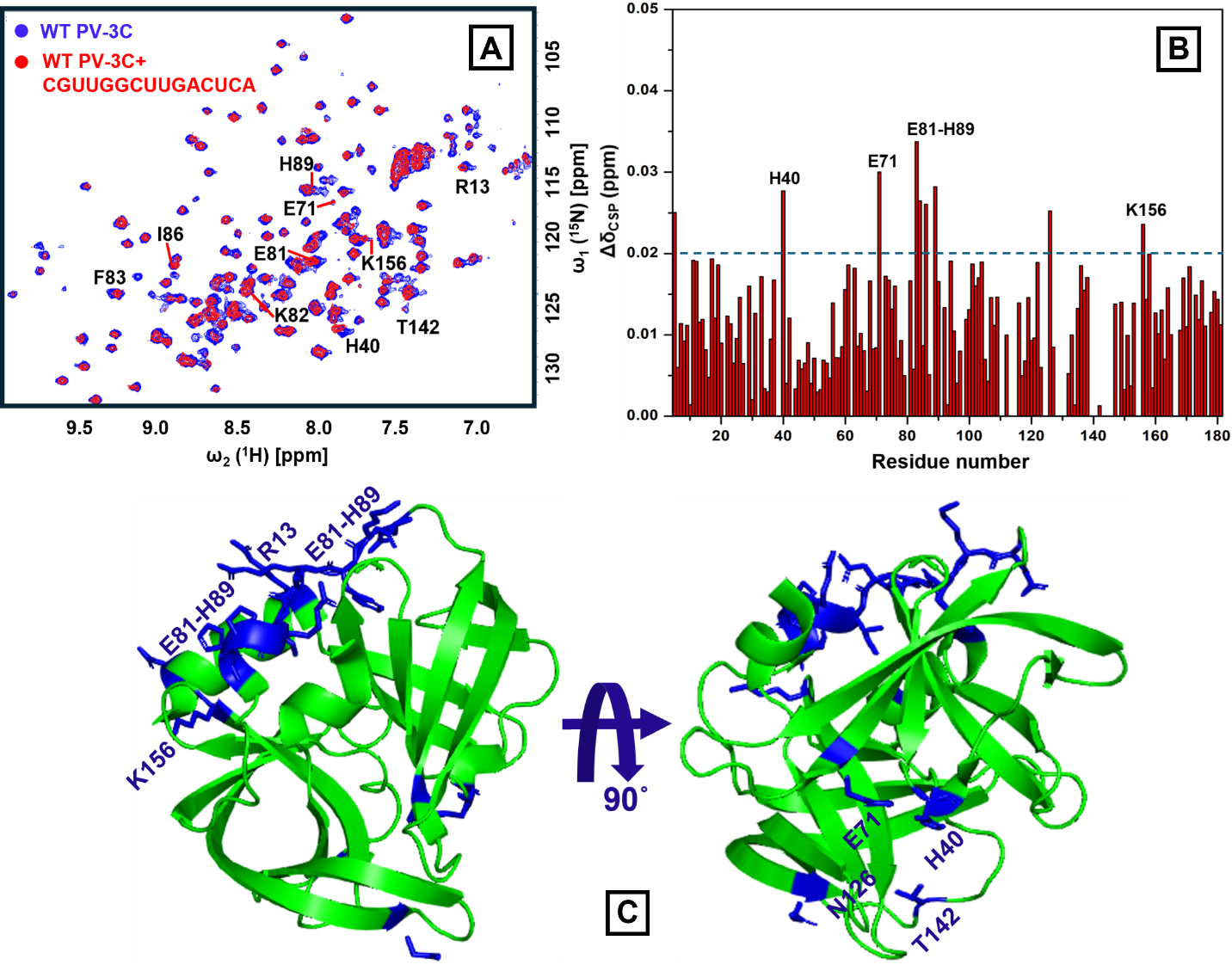

**Figure S8. PV-3C and RNA-9 (CGUUGGCUUGACUCA) binding.** (A) ^1^H-^15^N SOFAST-HMQC spectra were compared with free wild-type PV-3C (blue) and RNA-9 bound PV-3C (red) in 1:0.5 stoichiometry; the residues with substantial CSP are labelled in the spectra. (B) NMR CSP values for the amino acid residues of PV-3C are plotted, using the equation in Fig. S2. (C) Residues showing substantial CSPs in the presence of RNA are represented in blue in the 3C X-ray crystal structure (PDB ID: 1L1N) using PyMOL These residues include cluster-I and cluster-II residues as determined using RNA-1 (see Figure 1). In addition, N126, near cluster-II, also shows CSP. Sample and NMR spectrometer conditions were as in Fig. S2.

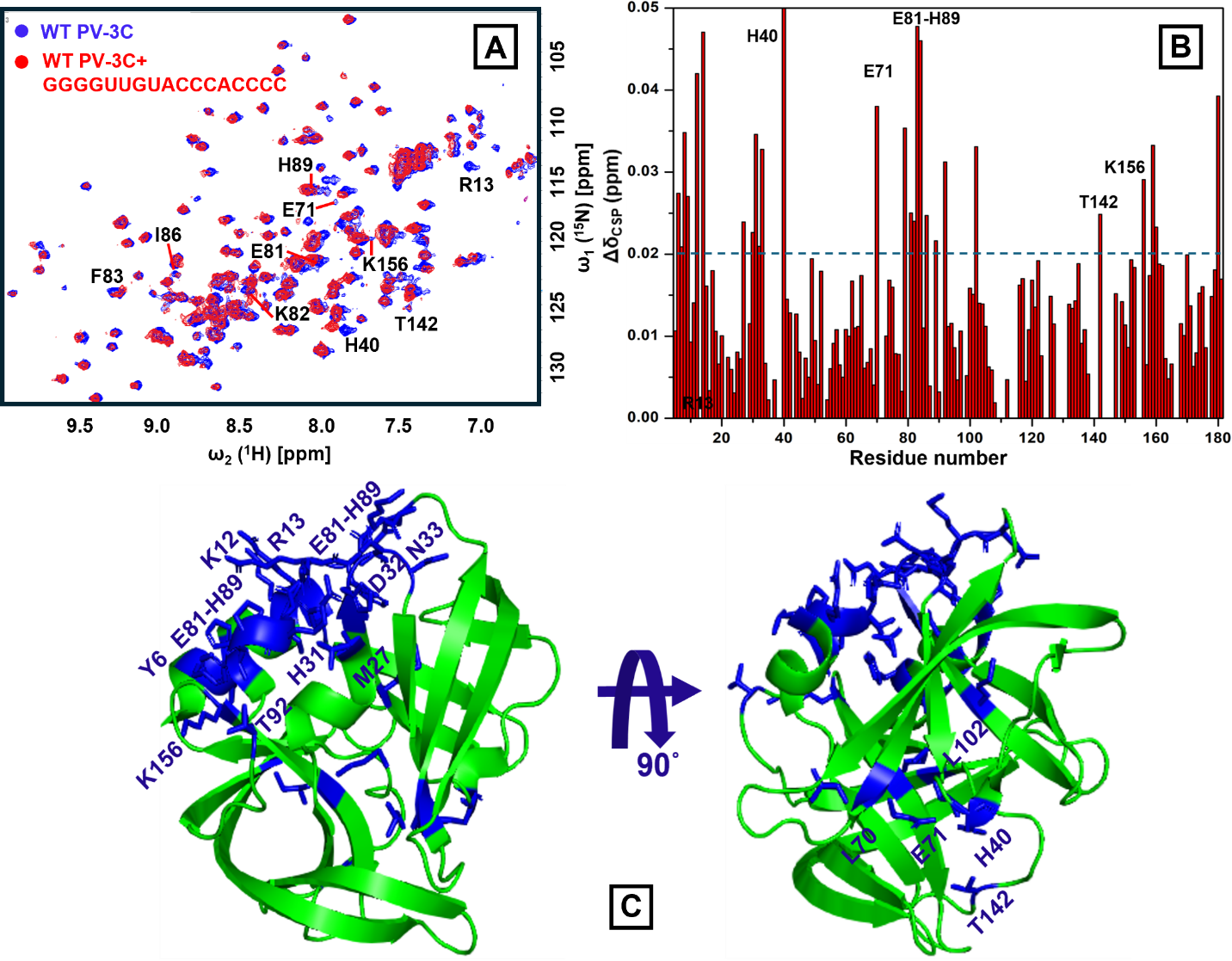

**Figure S9. PV-3C and RNA-10 (GGGGUUGUACCCACCCC) binding.** (A) ^1^H-^15^N SOFAST-HMQC spectra were compared with free wild-type PV-3C (blue) and RNA-10 bound PV-3C (red) in 1:0.5 stoichiometry; the residues with substantial CSP are labelled in the spectra. (B) NMR CSP values for the amino acid residues of PV-3C are plotted, using the equation in Fig. S2. (C) Residues showing substantial CSPs in the presence of RNA are represented in blue in the 3C X-ray crystal structure (PDB ID: 1L1N) using PyMOL These residues include cluster-I and cluster-II residues as determined using RNA-1 (see Figure 1). In addition, N14, M27, V30, H31, D32, N33, and T92, near cluster-I, and L70, and L102, near cluster-II, also show CSP. Sample and NMR spectrometer conditions were as in Fig. S2.

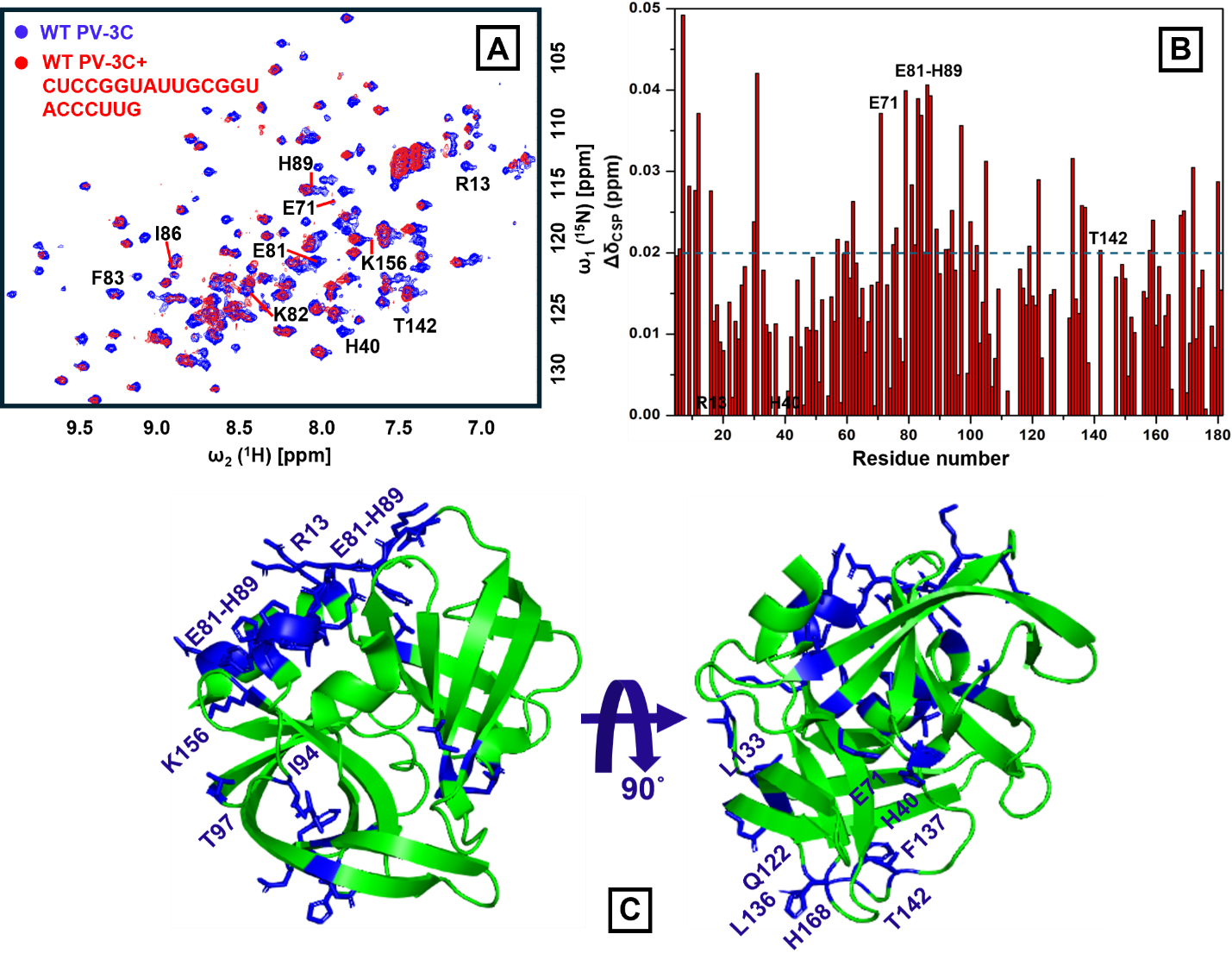

**Figure S10. PV-3C and RNA-11 (CUCCGGUAUUGCGGUACCCUUG) binding.** (A) ^1^H-^15^N SOFAST-HMQC spectra were compared with free wild-type PV-3C (blue) and RNA-11 bound PV-3C (red) in 1:0.5 stoichiometry; the residues with substantial CSP are labelled in the spectra. (B) NMR CSP values for the amino acid residues of PV-3C are plotted, using the equation in Fig. S2. (C) Residues showing substantial CSPs in the presence of RNA are represented in blue in the 3C X-ray crystal structure (PDB ID: 1L1N) using PyMOL These residues include cluster-I and cluster-II residues as determined using RNA-1 (see Figure 1). In addition, I94 and T97, near cluster-I and Q122, L133, L136, F137, and H168, near cluster-II, also show CSP. Sample and NMR spectrometer conditions were as in Fig. S2.

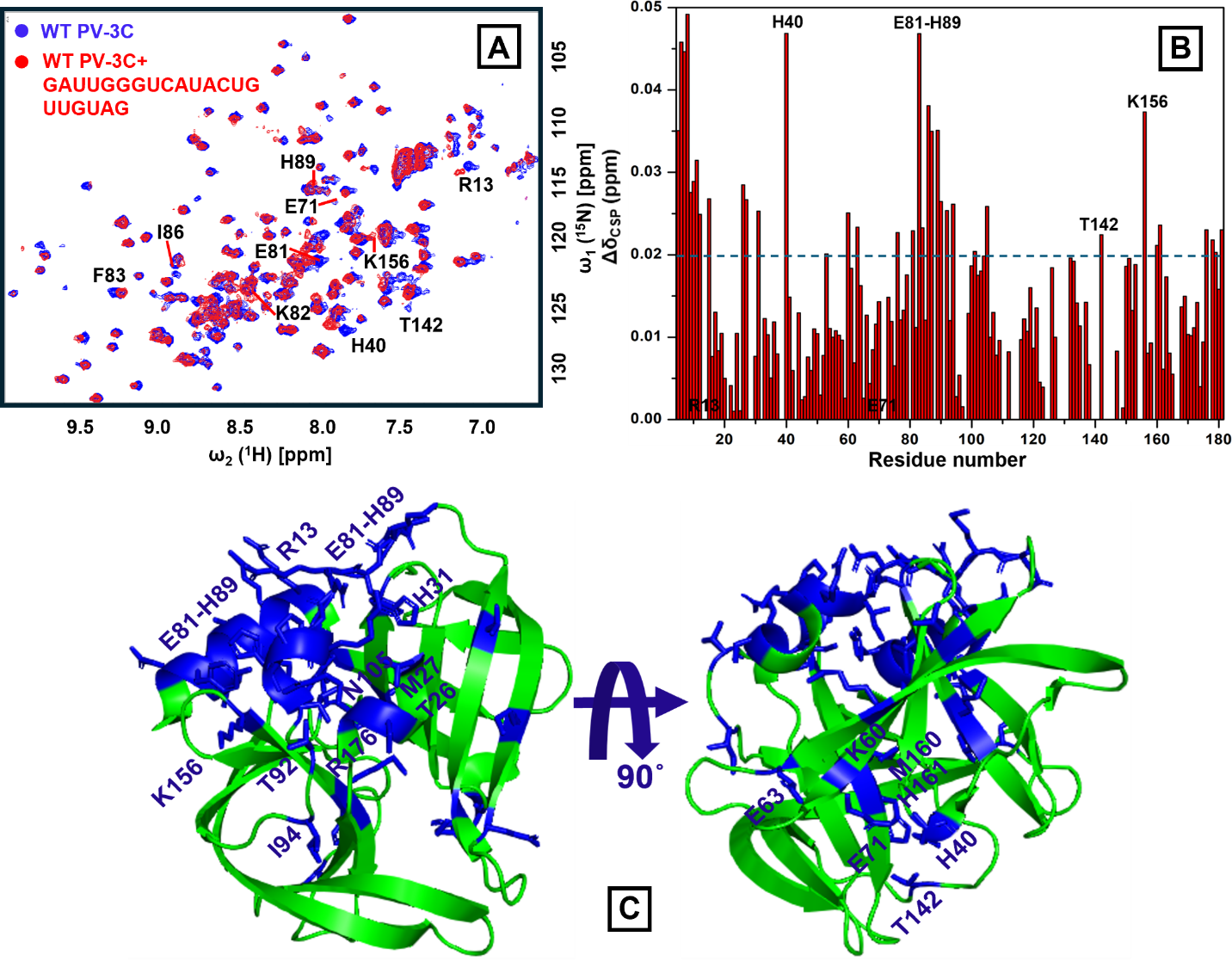

**Figure S11. PV-3C and RNA-12 (GAUUGGGUCAUACUGUUGUAG) binding.** (A) ^1^H-^15^N SOFAST-HMQC spectra were compared with free wild-type PV-3C (blue) and RNA-12 bound PV-3C (red) in 1:0.5 stoichiometry; the residues with substantial CSP are labelled in the spectra. (B) NMR CSP values for the amino acid residues of PV-3C are plotted, using the equation in Fig. S2. (C) Residues showing substantial CSPs in the presence of RNA are represented in blue in the 3C X-ray crystal structure (PDB ID: 1L1N) using PyMOL These residues include cluster-I and cluster-II residues as determined using RNA-1 (see Figure 1). In addition, T26, M27, H31, I90, T92, I94, N105, and R176, near cluster-I and K60, E63, M160, and H161, near cluster-II, also show CSP. Sample and NMR spectrometer conditions were as in Fig. S2.

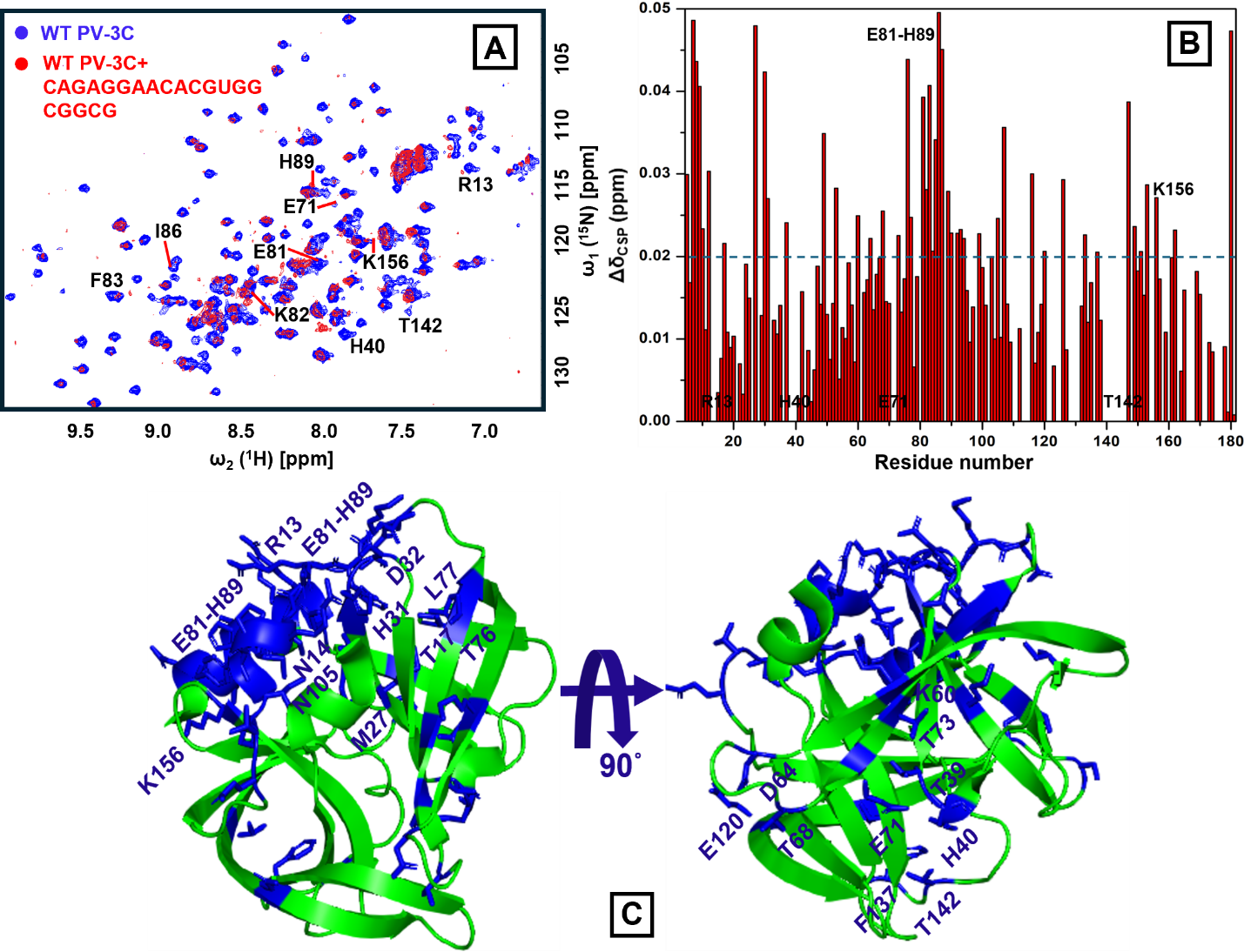

**Figure S12. PV-3C and RNA-13 (CAGAGGAACACGUGGCGGCG) binding.** (A) ^1^H-^15^N SOFAST-HMQC spectra were compared with free wild-type PV-3C (blue) and RNA-13 bound PV-3C (red) in 1:0.5 stoichiometry; the residues with substantial CSP are labelled in the spectra. (B) NMR CSP values for the amino acid residues of PV-3C are plotted, using the equation in Fig. S2. (C) Residues showing substantial CSPs in the presence of RNA are represented in blue in the 3C X-ray crystal structure (PDB ID: 1L1N) using PyMOL These residues include cluster-I and cluster-II residues as determined using RNA-1 (see Figure 1). In addition, N14, T17, M27, V30, H31, D32, T76, L77, and N105, near cluster-I and T39, K60, D64, T68, T73, E120, and F137, near cluster-II, also show CSP. Sample and NMR spectrometer conditions were as in Fig. S2.

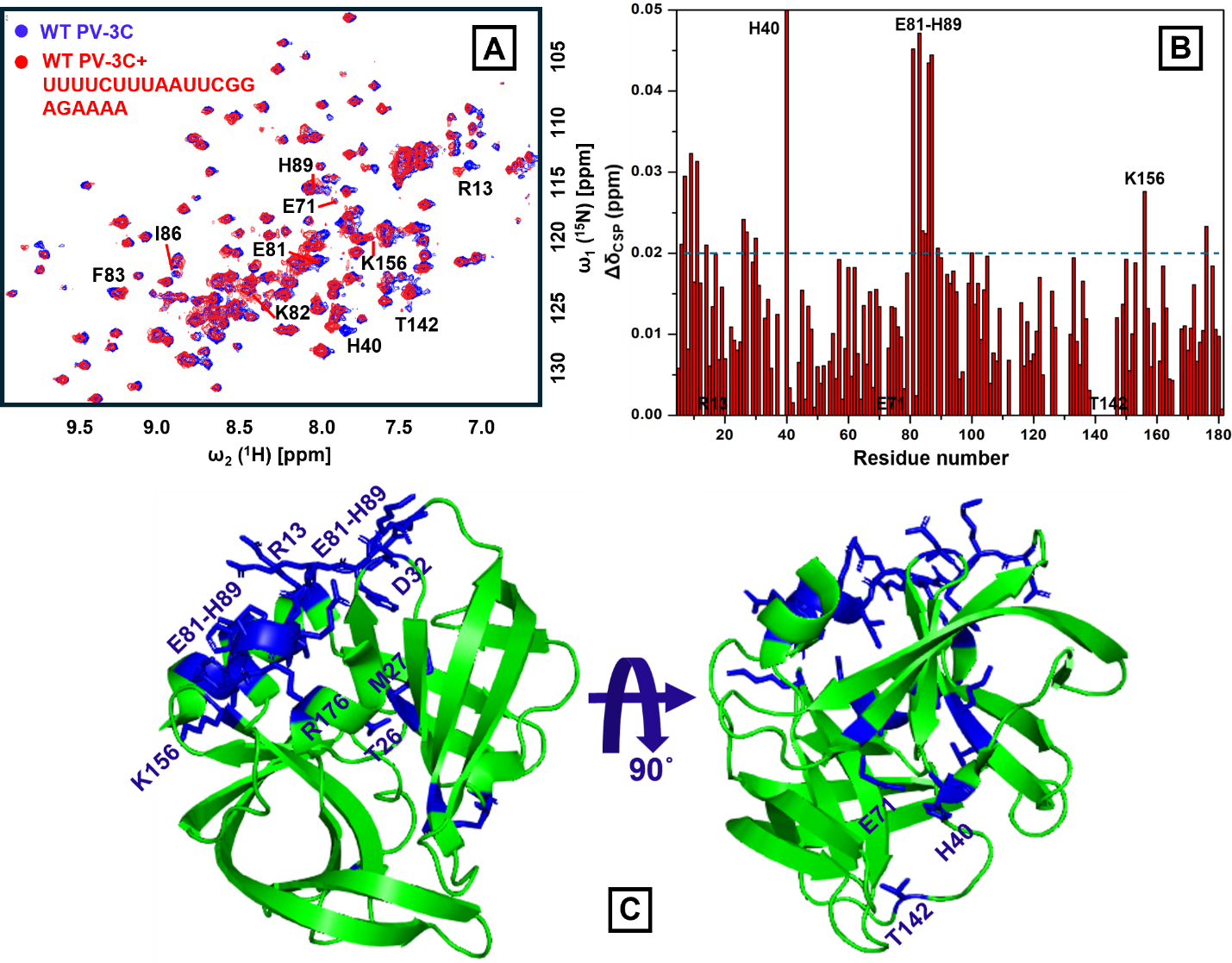

**Figure S13. PV-3C and RNA-14 (UUUUCUUUAAUUCGGAGAAAA) binding.** (A) ^1^H-^15^N SOFAST-HMQC spectra were compared with free wild-type PV-3C (blue) and RNA-14 bound PV-3C (red) in 1:0.5 stoichiometry; the residues with substantial CSP are labelled in the spectra. (B) NMR CSP values for the amino acid residues of PV-3C are plotted, using the equation in Fig. S2. (C) Residues showing substantial CSPs in the presence of RNA are represented in blue in the 3C X-ray crystal structure (PDB ID: 1L1N) using PyMOL These residues include cluster-I and cluster-II residues as determined using RNA-1 (see Figure 1). In addition, T26, M27, D32, and R176, near cluster-I, also show CSP. Sample and NMR spectrometer conditions were as in Fig. S2.

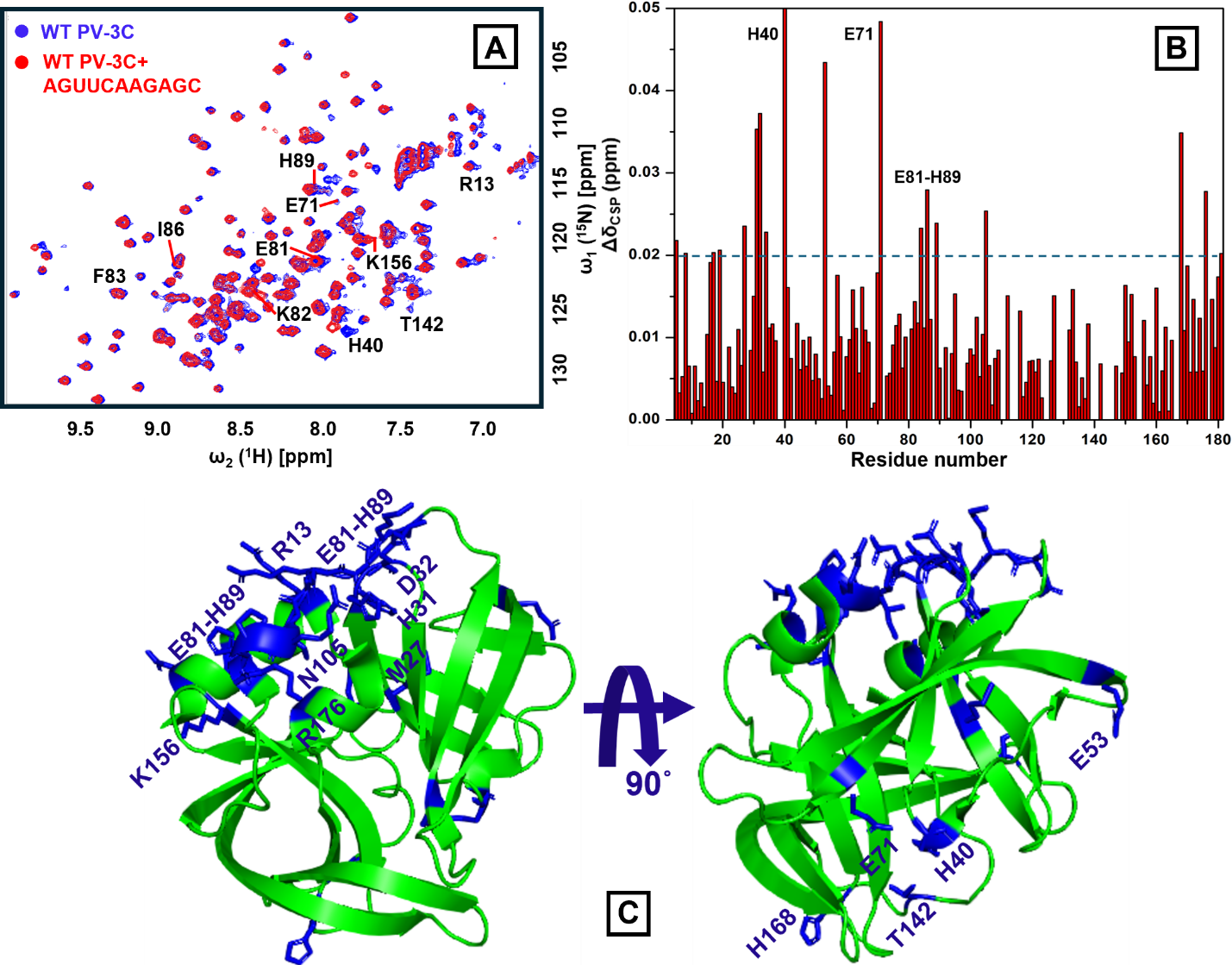

**Figure S14. PV-3C and RNA-15 (AGUUCAAGAGC) binding.** (A) ^1^H-^15^N SOFAST-HMQC spectra were compared with free wild-type PV-3C (blue) and RNA-15 bound PV-3C (red) in 1:0.5 stoichiometry; the residues with substantial CSP are labelled in the spectra. (B) NMR CSP values for the amino acid residues of PV-3C are plotted, using the equation in Fig. S2. (C) Residues showing substantial CSPs in the presence of RNA are represented in blue in the 3C X-ray crystal structure (PDB ID: 1L1N) using PyMOL These residues include cluster-I and cluster-II residues as determined using RNA-1 (see Figure 1). In addition, M27, H31, D32, N105, and R176, near cluster-I, and E53, and H168, near cluster-II, also show CSP. Sample and NMR spectrometer conditions were as in Fig. S2.

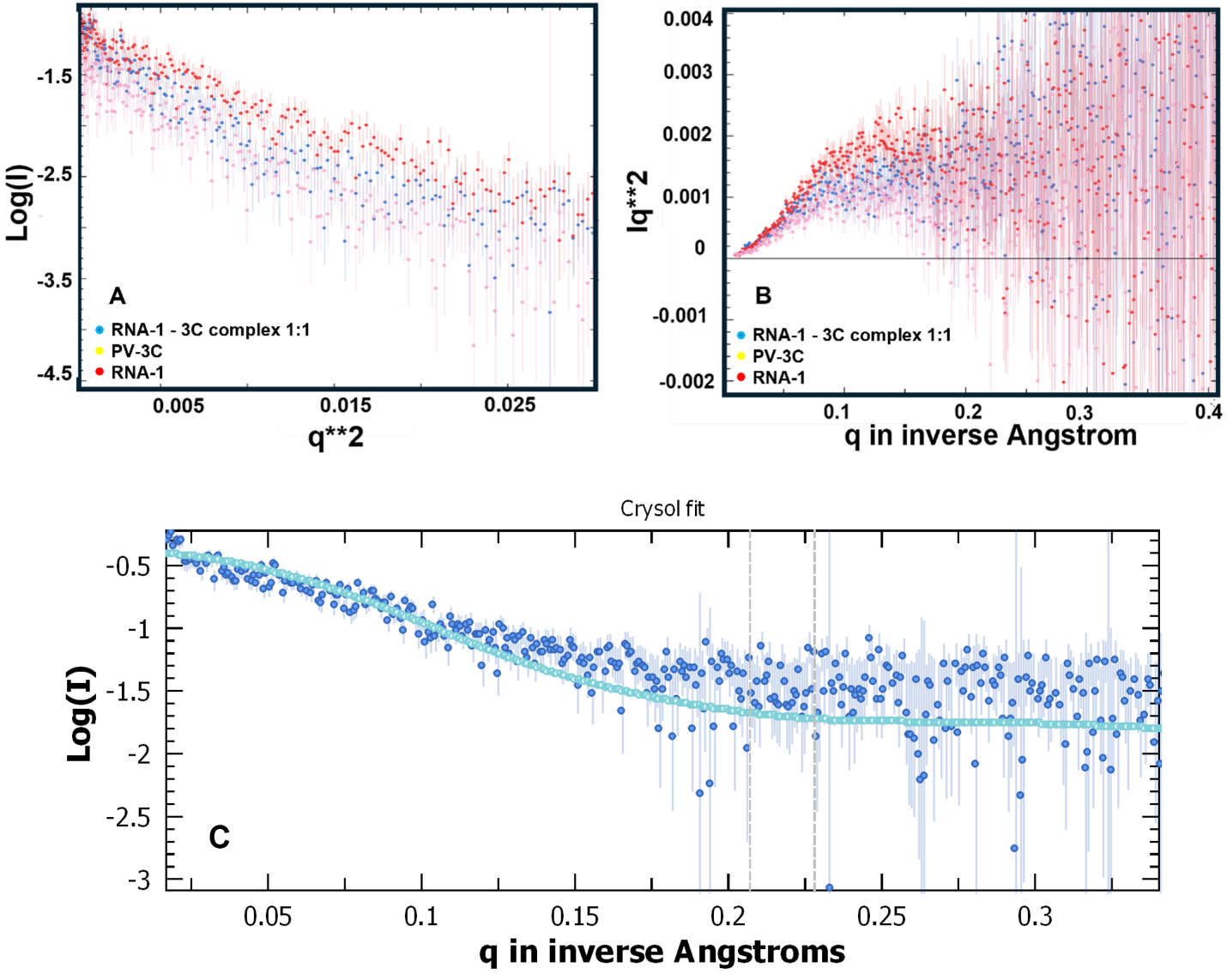

**Figure S15.** **Guinear data overlay, Kratky overlay and Crysol fit for 3C, RNA-1 and RNA-1 - 3C complex (1:1) from SAXS measurement.** (A) Guinear plot of Log (I) vs q**2 showing linear region indicating the particle is monodisperse (uniform in size) and globular. (B) Kratky plot of Iq**2 vs q in inverse Angstrom are represented here to understand the conformational change, (C) Crysol fit of Log(I) vs q in inverse Angstroms to overlay the computed scattering curve as from the model to the experimental SAXS scattering profile

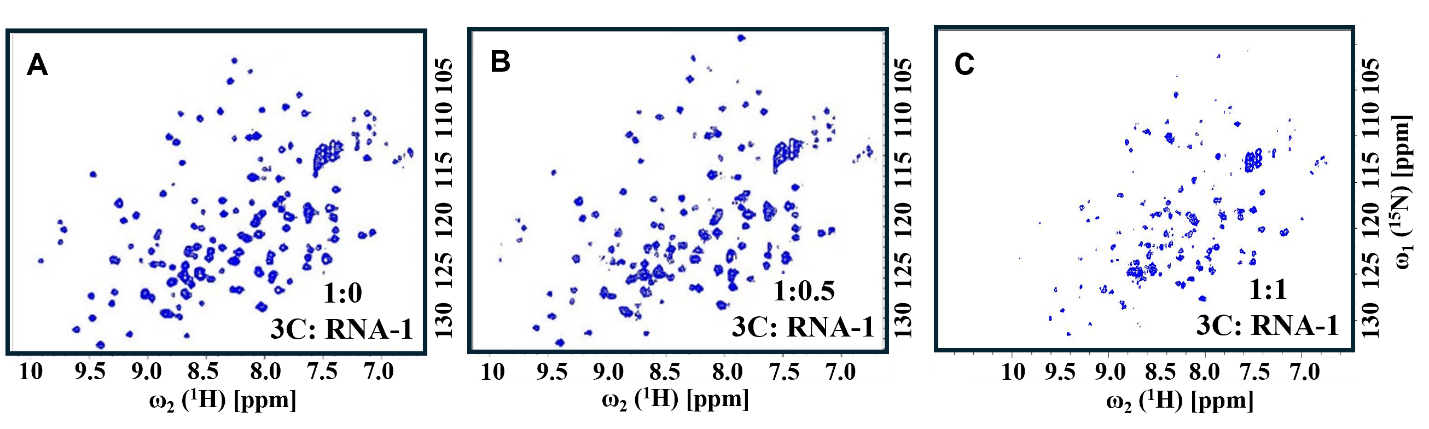

**Figure S16.** **Disappearance of NMR peaks at higher ratios of 3C: RNA-1 suggests multimeric complex formation.** 2D [^15^N-^1^H] SOFAST-HMQC NMR spectra of free 3C (A), after the addition of 1:0.5 (B) and 1:1 (C) stoichiometry addition of 3C: RNA-1 respectively. The PV-3C protein concentration was 25 µM and RNA was added following a 1:0.5 and 1:1 stoichiometry. Both protein and RNA were in a buffer containing 10 mM HEPES, 50 mM NaCl, and pH 7.5. NMR spectrometer conditions were as in Fig. S2.

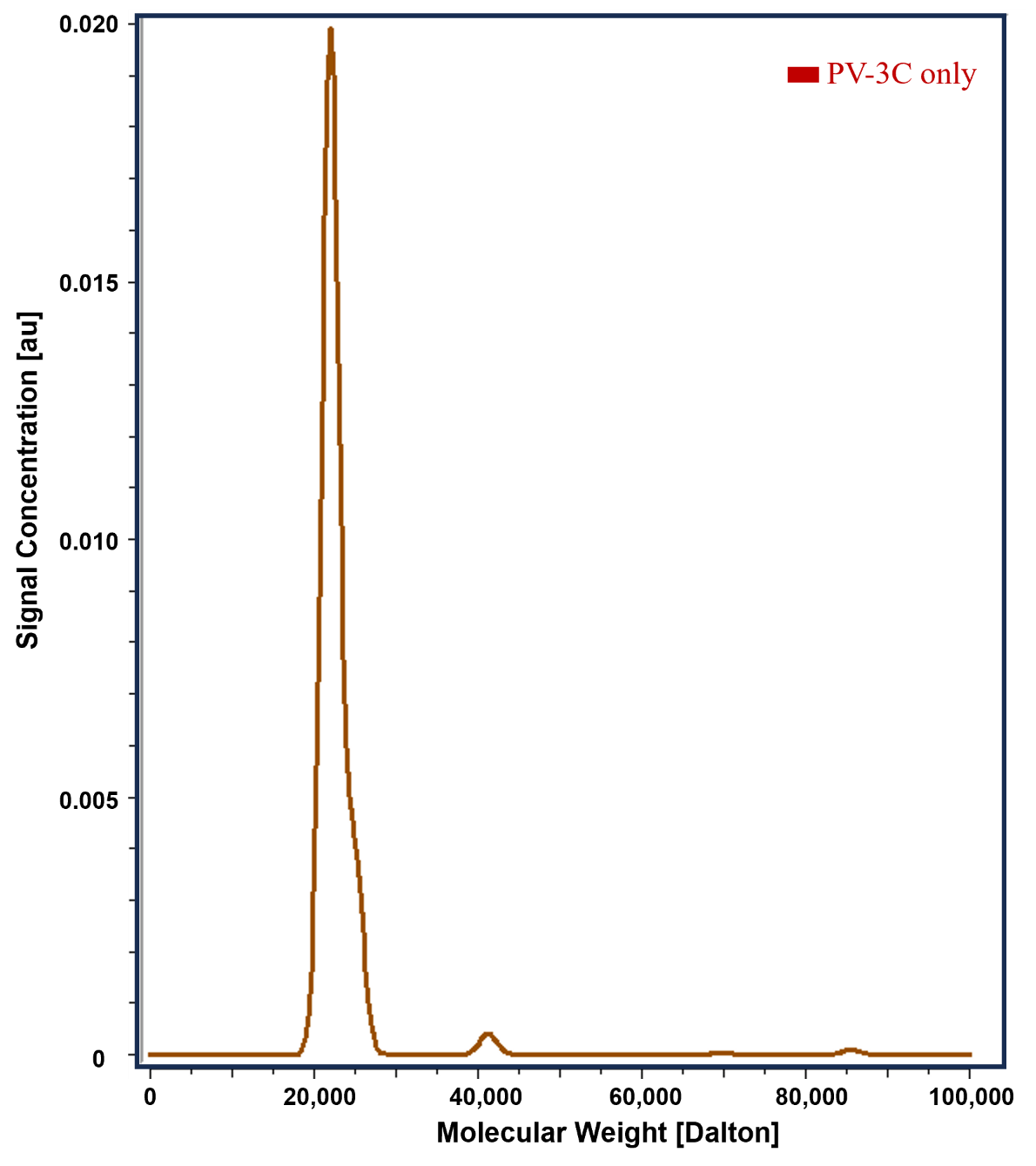

**Figure S17. SV-AUC of PV-3C only.** PV-3C is predominantly in monomeric form at a concentration of 50 µM. All our biophysical experiments were performed below this protein concentration. PV-3C was in a buffer containing 10 mM HEPES, 50 mM NaCl, and pH 7.5.

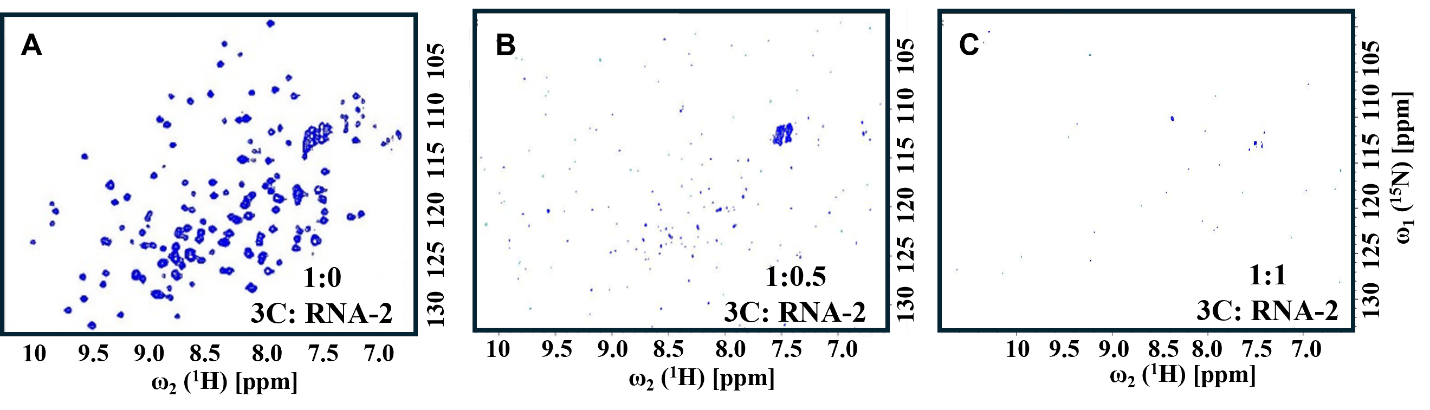

**Figure S18.** **Titration with RNA-2 leads to disappearance of NMR peaks at even lower** **3C to RNA ratios.** 2D [^15^N-^1^H] SOFAST-HMQC NMR spectra of free 3C (A), after the addition of 1:0.5 (B) and 1:1 (C) stoichiometry addition of 3C: RNA-2 respectively. The PV-3C protein concentration was 25 µM and RNA was added following a 1:0.5 and 1:1 stoichiometry. Both protein and RNA were in a buffer containing 10 mM HEPES, 50 mM NaCl, and pH 7.5.

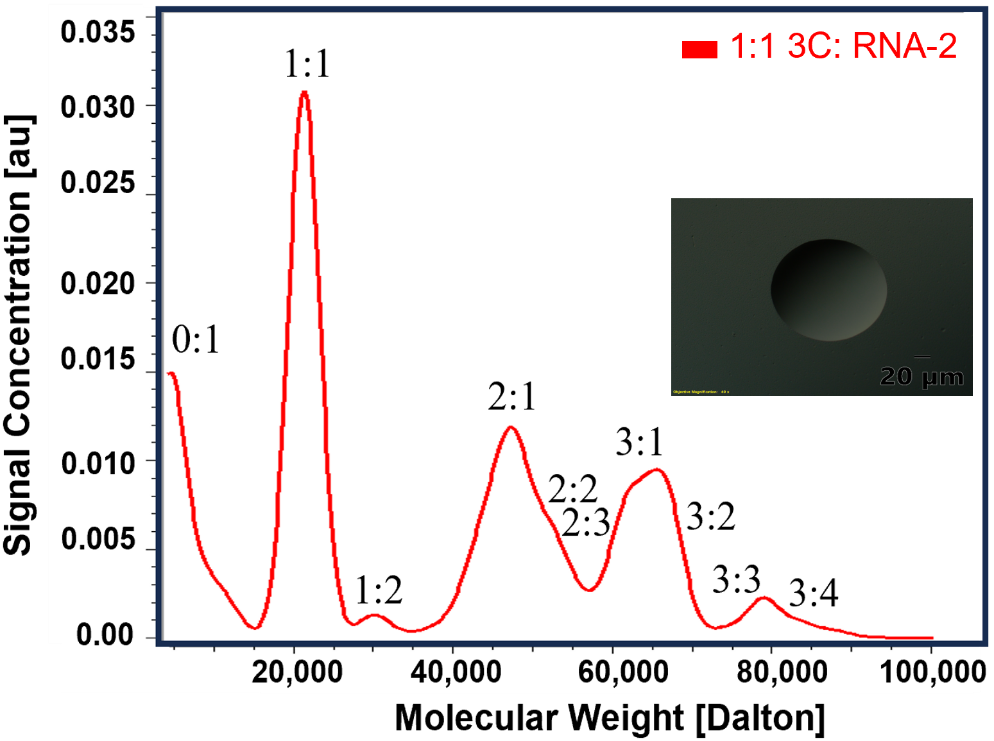

**Figure S19. Multimeric complex formation between PV-3C and RNA-2 leads to LLPS formation.** PV-3C and RNA-2 form various multimolar-ratio complexes at 1:1 molar stoichiometry, which results in LLPS observed through a DIC microscope as shown in the inset. The PV-3C protein concentration was 25 µM and RNA was added following a 1:1 stoichiometry. Both protein and RNA were in a buffer containing 10 mM HEPES, 50 mM NaCl, and pH 7.5.

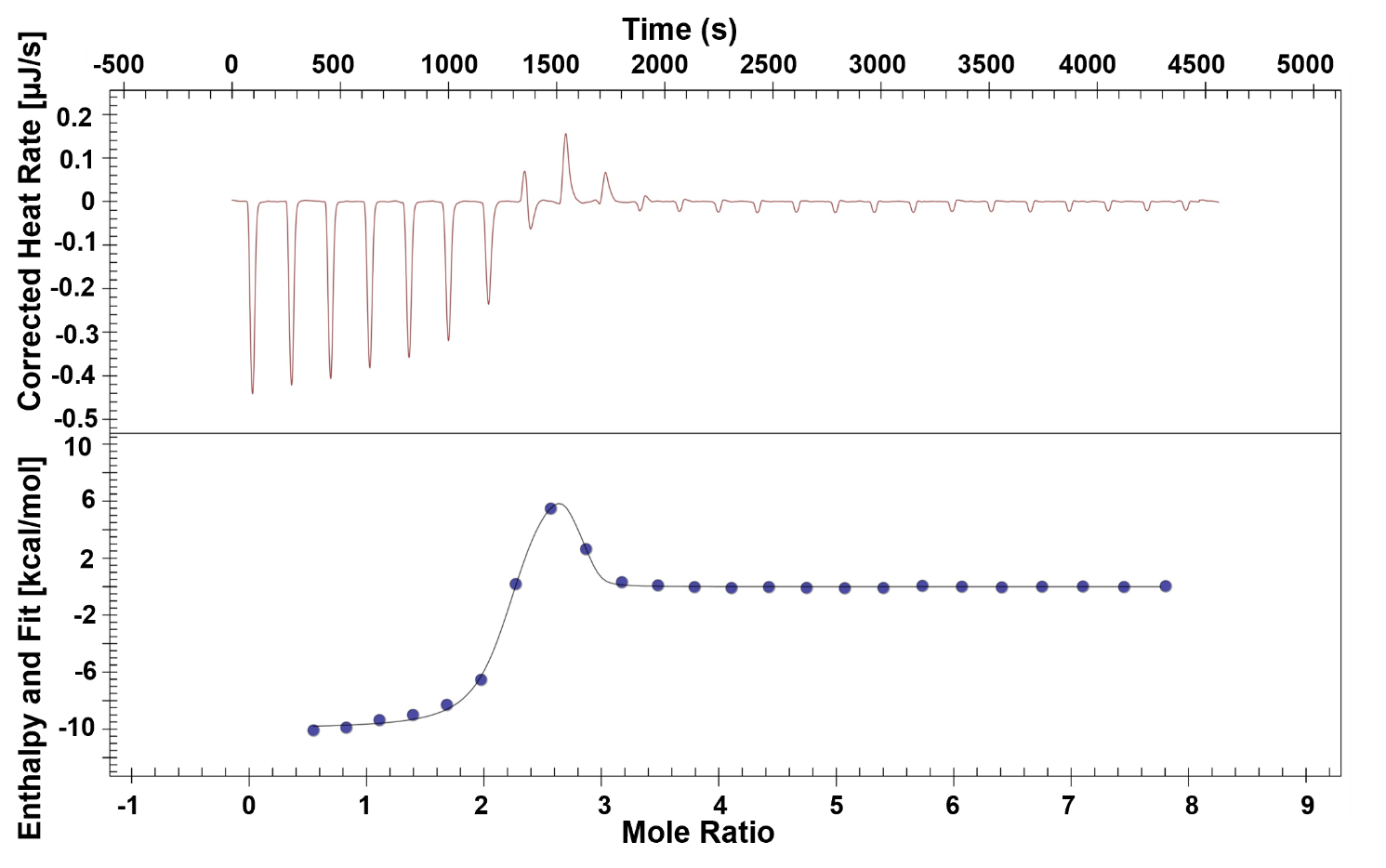

**Figure S20. ITC suggests two RNA binding sites on PV-3C.** Two binding sites on PV-3C for RNA-1 were demonstrated using ITC, with 3C at a concentration of 400 µM titrated into RNA at 20 µM. A second titration set was carried out with 400 µM of 3C into the buffer (10 mM HEPES, pH 7.5, and 50 mM NaCl) for baseline subtraction. Each experiment involved 25 injections of 2.0 µL each.

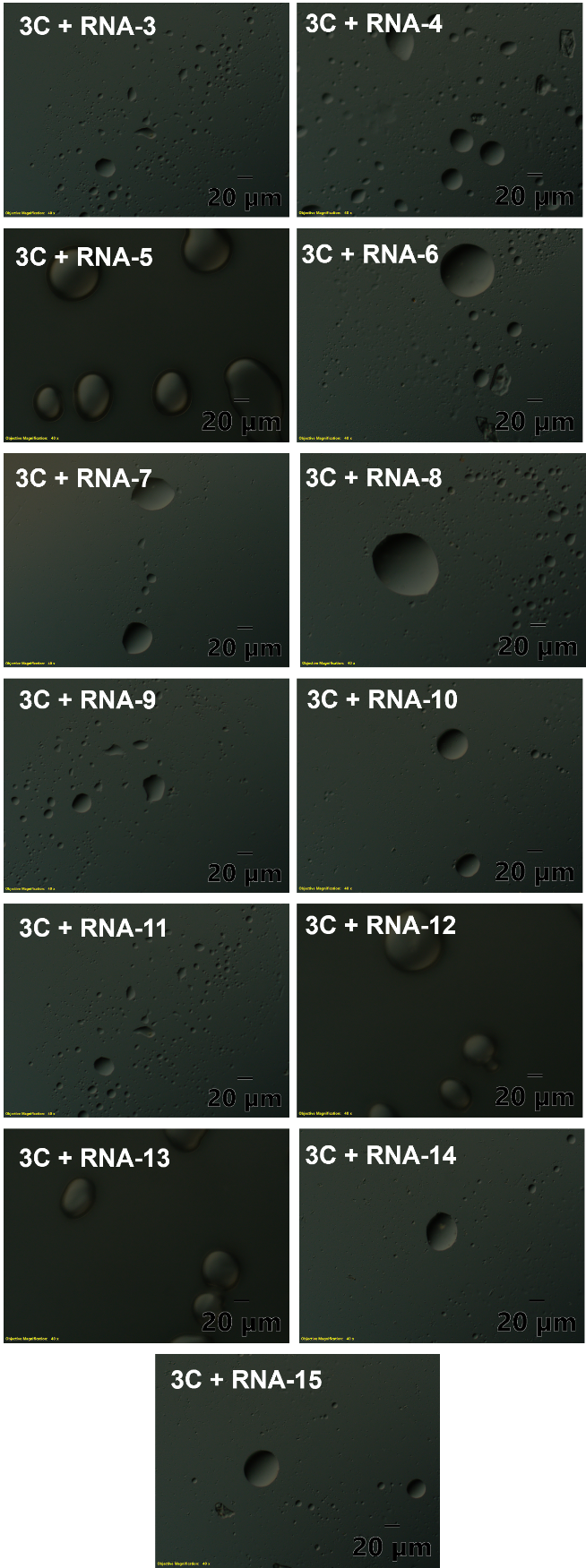

**Figure S21. LLPS formation of PV-3C with RNA-3- RNA-15 observed through DIC microscopy.**

DIC microscopy shows the LLPS formation of PV-3C with the RNA-3- RNA-15 individually. PV-3C samples with RNA in the HEPES buffer (10 mM HEPES pH 7.5 and 50 mM NaCl) were prepared, and a drop was mounted on clean glass slides with a coverslip for high-quality DIC microscopy imaging. There was no staining, and appropriate control of buffer, protein, and RNA were recorded before observing LLPS in the protein-RNA samples.

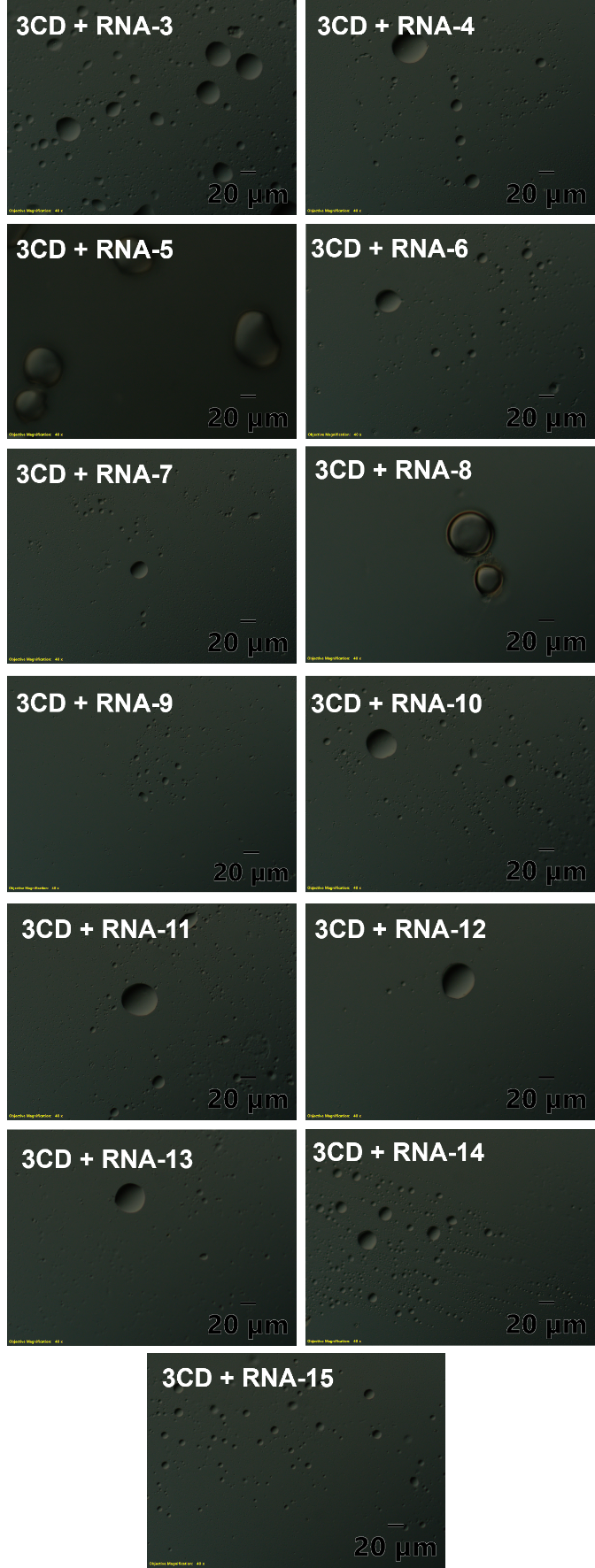

**Figure S22. LLPS formation of PV-3CD with RNA-3- RNA-15 observed through DIC microscopy.**

DIC microscopy shows the LLPS formation of PV-3CD with the RNA-3- RNA-15 individually.PV-3CD samples with RNA in the HEPES buffer (10 mM HEPES pH 7.5 and 50 mM NaCl) were prepared, and a drop was mounted on clean glass slides with a coverslip for high-quality DIC microscopy imaging. There was no staining, and appropriate control of buffer, protein, and RNA were recorded before observing LLPS in the protein-RNA samples.

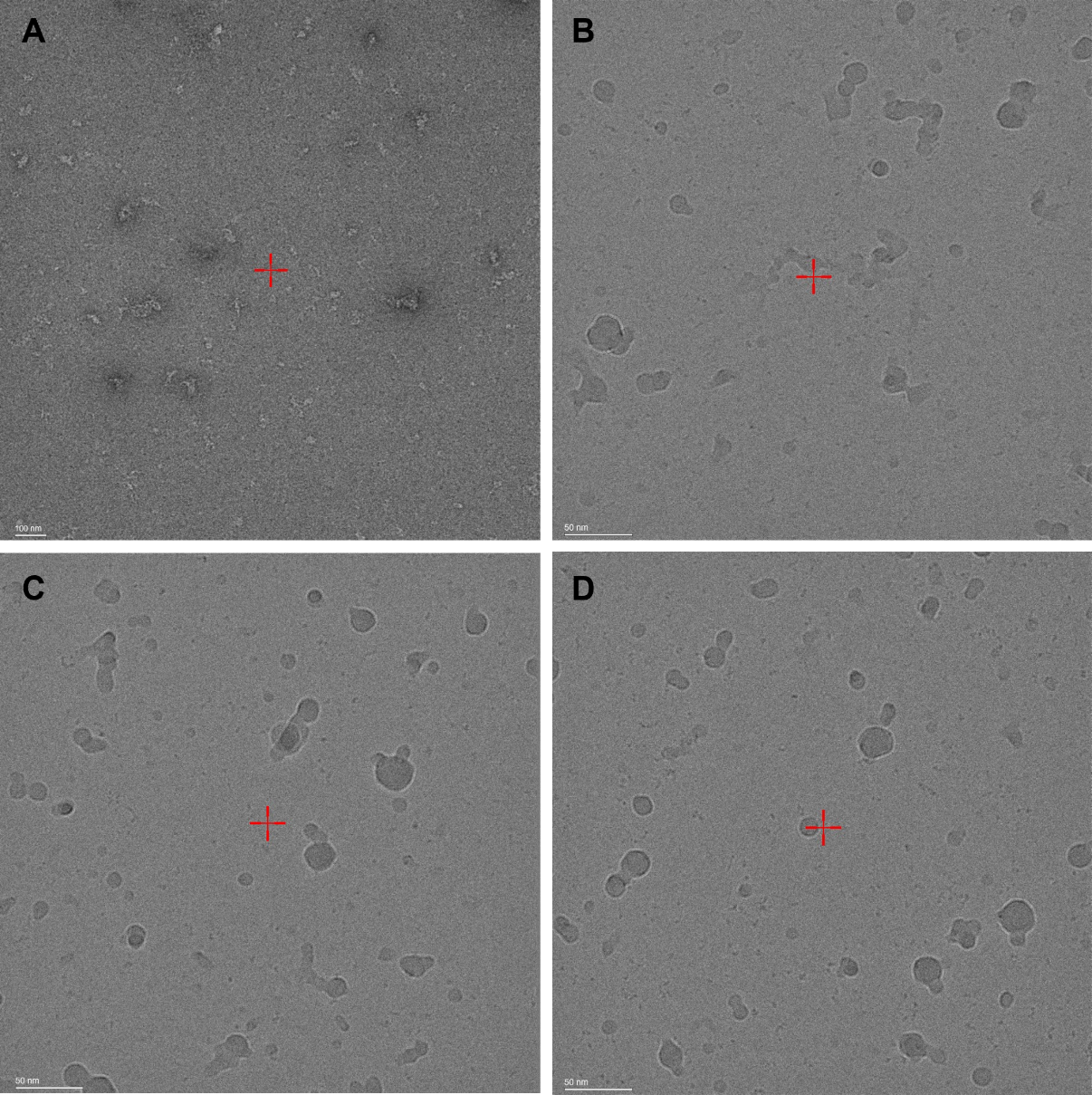

**Figure S23.** **EM analysis of 3CD-RNA-1 oligomerization.** Representative EM micrographs displaying the formation of oligomers in the range of ~20-40 nm. A 3.5 μL aliquot of the sample was applied to a carbon support TEM grid (400 mesh, Ted Pella) and incubated for 1 minute. Subsequently, excess liquid was carefully blotted away using filter paper. The grid was then washed twice with 10 μL of ultrapure water to remove any unbound material. The grid was then stained with 10 μL of freshly prepared uranyl formate solution (0.7% w/v, pH 4.5) for 30 seconds. Finally, after blotting away the excess staining solution, the grid was air-dried at room temperature. The details of the negative staining electron microscopy are described in the ‘Method’ section.

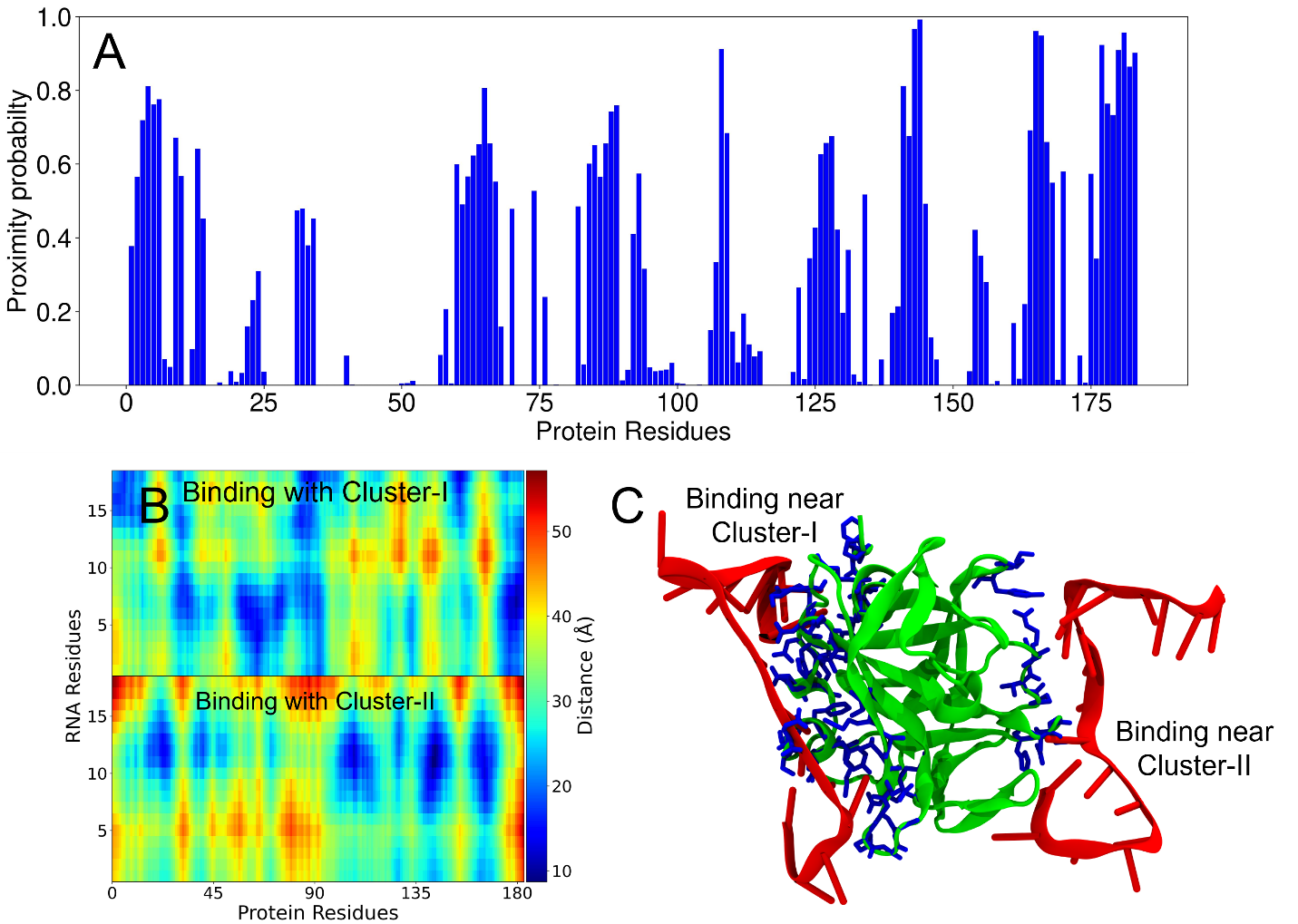

**Figure S24. Interactions between PV-3C protein and RNA-2 from MD simulations.** (A) Proximity probabilities (*P_prox_*) of residues in 3C. (B) Distance map between the residues of RNA-2 and 3C. The color bar represents the color-coded distances underlying the distance map. The vertical axis shows the RNA residues of the two chains (each with 18 nucleotides). (C) Simulation snapshot showing two chains of RNA-2 interacting with the residues in Cluster-I and Cluster-II of the 3C protein (as also observed from NMR experiments). The protein and RNA chains are shown in green and red respectively. The residues colored blue have proximity probabilities greater than 0.5.

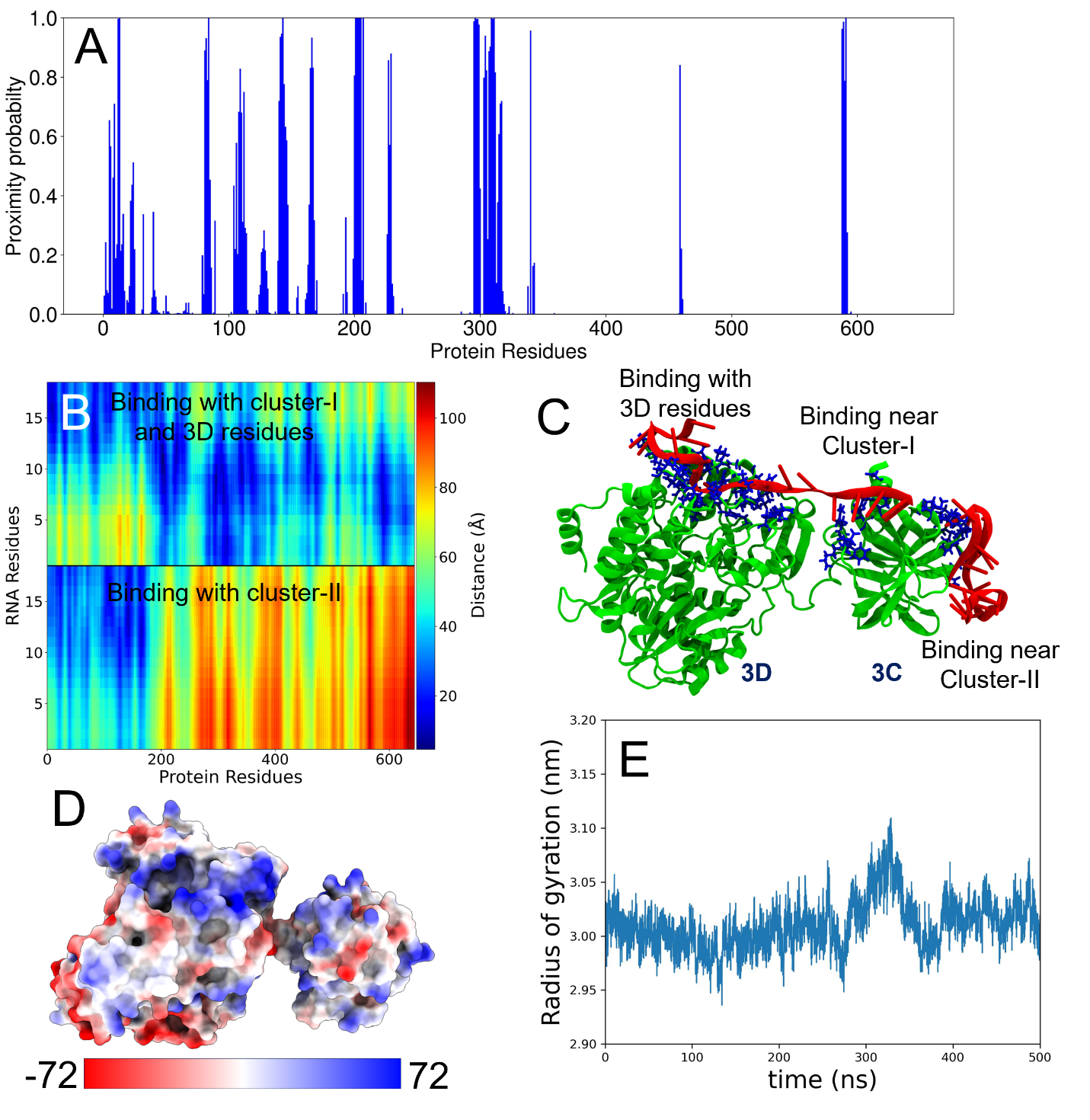

**Figure S25. Interactions between PV-3CD protein and RNA-2 from MD simulations.** (A) Proximity probabilities (*P_prox_*) of residues in 3CD. (B) Distance map between the residues of RNA-2 and 3CD. The color bar represents the color-coded distances underlying the distance map. The vertical axis shows the RNA residues of the two chains (each with 18 nucleotides). (C) Simulation snapshot showing two chains of RNA-2 interacting with the residues in Cluster-I and Cluster-II of the 3CD protein. The former also interacts with positively charged residues in the 3D domain. The protein and RNA chains are shown in green and red respectively. The residues colored blue have proximity probabilities greater than 0.5. (D) Electrostatic map of the PV-3CD protein surface. (E) Time evolution of the radius of gyration (Rg) of PV-3CD obtained from MD simulations.

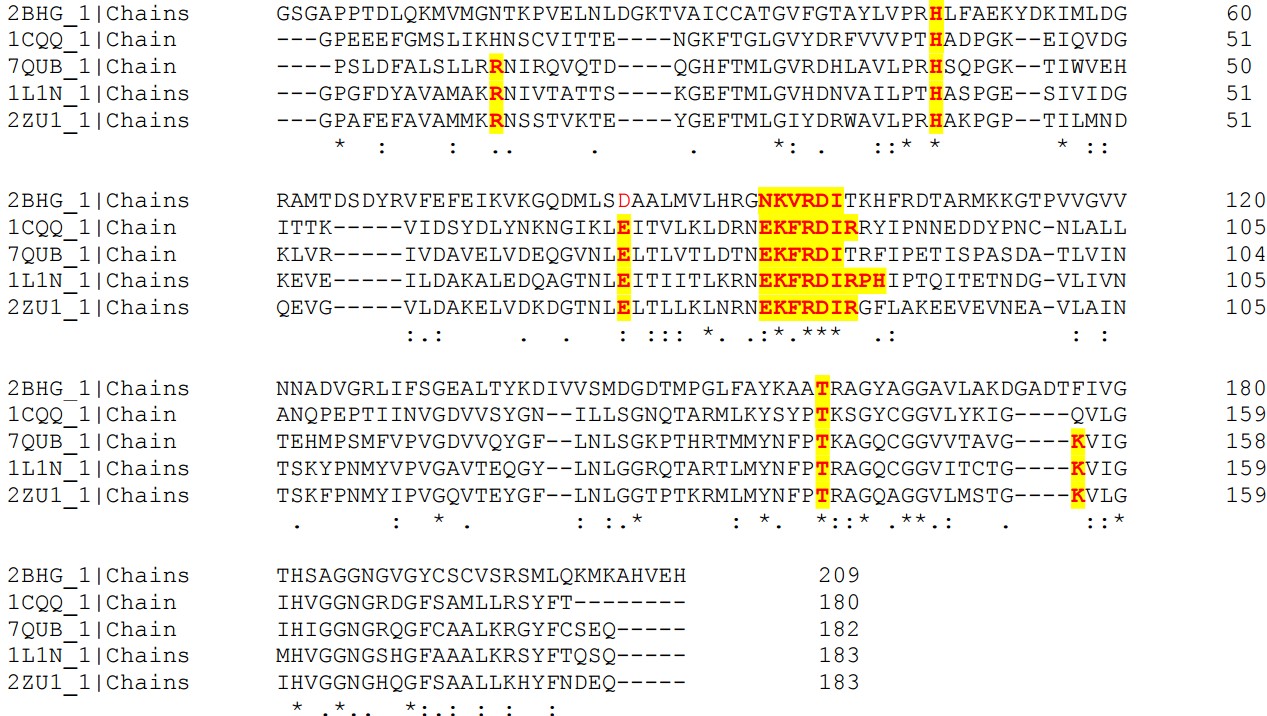

**Figure S26. CLUSTAL O (1.2.4) multiple sequence alignment of 3C proteases.** Clustal omega multiple sequence alignments of PV-3C (PDB ID: 1L1N) with Coxsackievirus B3 (CVB 3C, PDB ID: 2ZU1), Enterovirus A71 (EVA71 3C, PDB ID: 7QUB), Type 2 Rhinovirus 3C protease (HRV 3C, PDB ID: 1CQQ), and Foot and Mouth Disease Virus (PDB ID: 2BHG) are presented. RNA binding residues from Cluster-I and Cluster-II sequentially align in other 3C proteases and PV-3C.

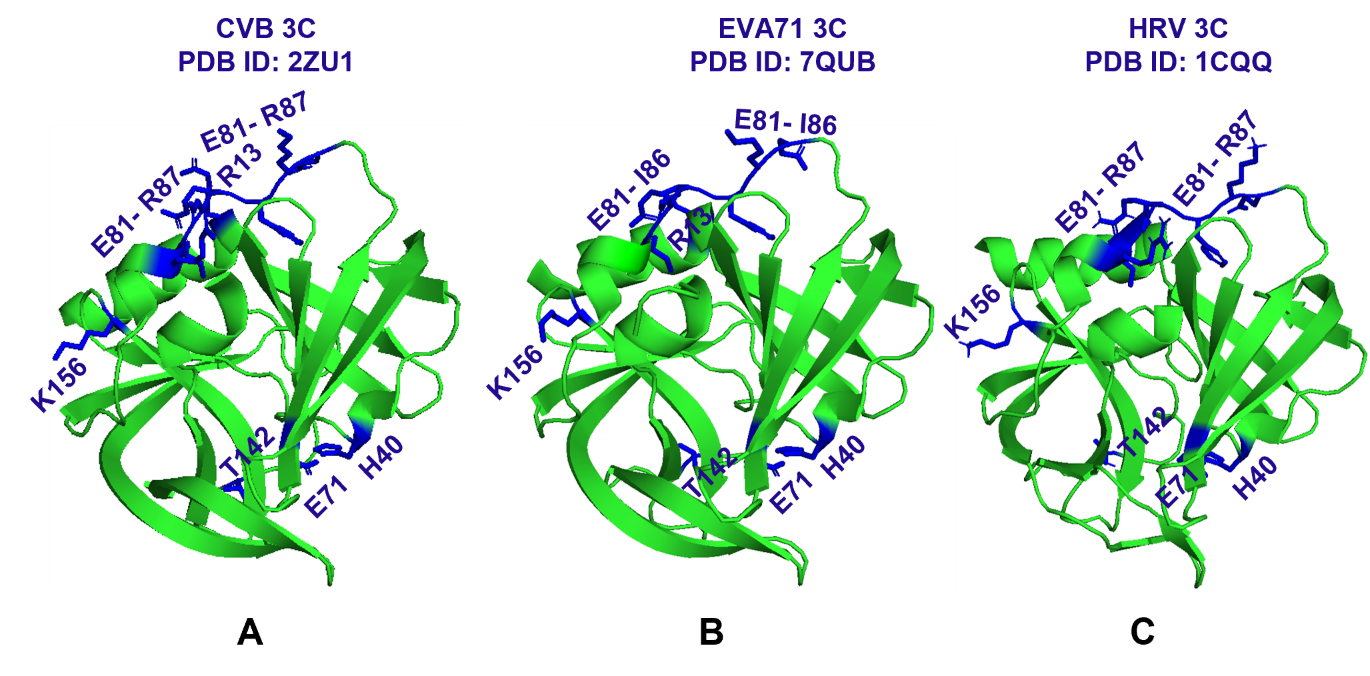

**Figure S27. The structural and sequential resemblance of PV-3C to other viral 3Cs.** PyMOL structures show the residues with high CSP obtained from PV-3C (Fig. 1) to be labelled on the structurally and sequentially resembled (A) 2ZU1 (CVB3 3C), (B) 7QUB (EV A71 3C) and (C) 1CQQ (HRV 3C). These three 3C proteases have similar sequential alignment to PV-3C **(Fig. S25)**.

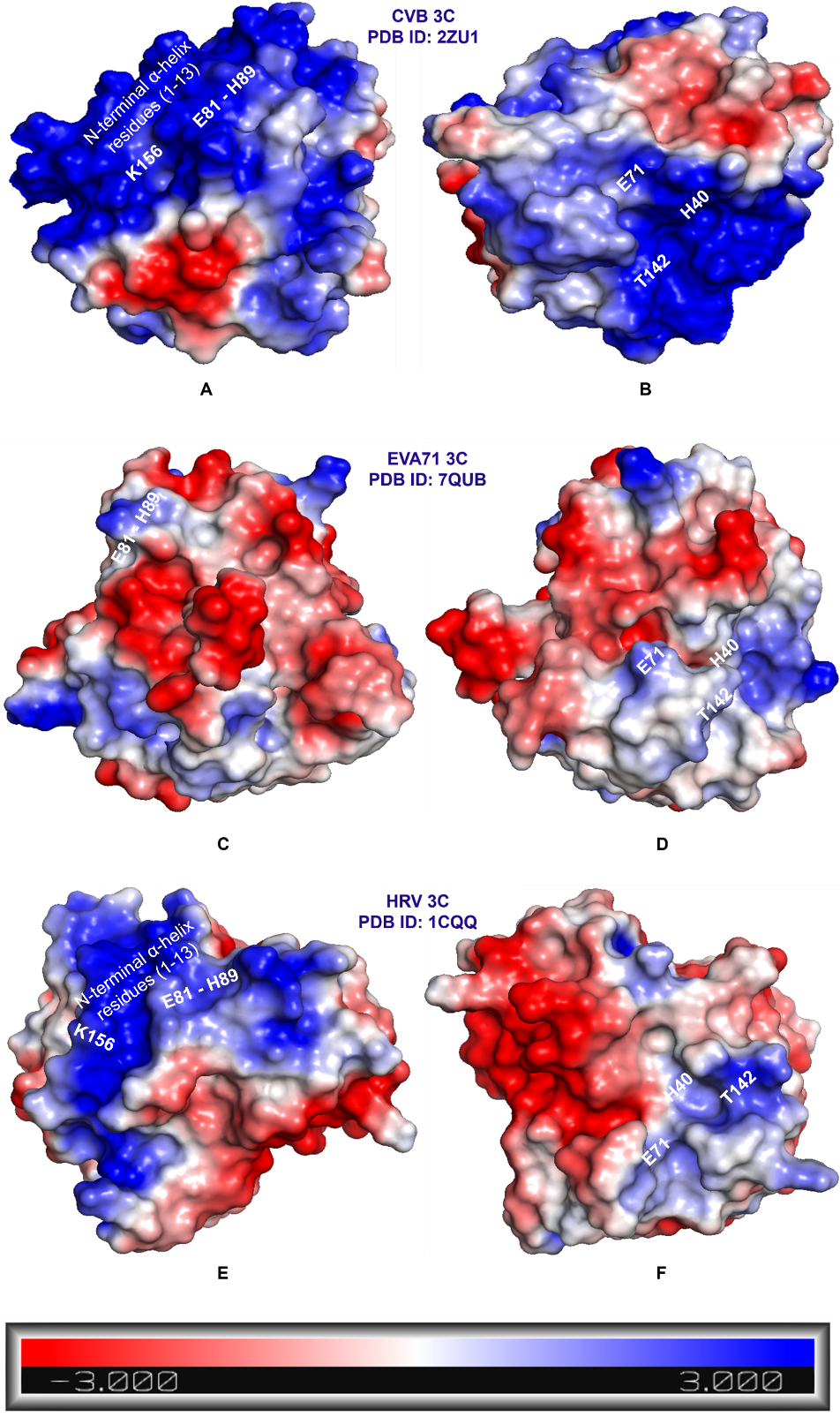

**Figure S28.** The electrostatic surface potential map for 3Cs, highlighting Cluster-I (A, C, E) and Cluster-II (B, D, F) regions, with blue and red representing positively- and negatively charged regions for CVB 3C, EV A71 3C, and HRV 3C respectively.
